# The Neurodata Without Borders ecosystem for neurophysiological data science

**DOI:** 10.1101/2021.03.13.435173

**Authors:** Oliver Rübel, Andrew Tritt, Ryan Ly, Benjamin K. Dichter, Satrajit Ghosh, Lawrence Niu, Ivan Soltesz, Karel Svoboda, Loren Frank, Kristofer E. Bouchard

## Abstract

The neurophysiology of cells and tissues are monitored electrophysiologically and optically in diverse experiments and species, ranging from flies to humans. Understanding the brain requires integration of data across this diversity, and thus these data must be findable, accessible, interoperable, and reusable (FAIR). This requires a standard language for data and metadata that can coevolve with neuroscience. We describe design and implementation principles for a language for neurophysiology data. Our open-source software (Neurodata Without Borders, NWB) defines and modularizes the interdependent, yet separable, components of a data language. We demonstrate NWB’s impact through unified description of neurophysiology data across diverse modalities and species. NWB exists in an ecosystem, which includes data management, analysis, visualization, and archive tools. Thus, the NWB data language enables reproduction, interchange, and reuse of diverse neurophysiology data. More broadly, the design principles of NWB are generally applicable to enhance discovery across biology through data FAIRness.

## Introduction

The immense diversity of life on Earth^1^ has always provided both inspiration and insight for biologists. For example, in neuroscience, the functioning of the brain is studied in species ranging from flies, to mice, to humans (**Fig. 1a**)^2^. Because brains evolved to produce a plethora of behaviors that advance organismal survival, neuroscientists monitor brain activity with a variety of different tasks and neural recording techniques. (**Fig. 1a**). These technologies provide complementary views of the brain, and creating a coherent model of how the brain works will require synthesizing data generated by these heterogeneous experiments. However, the extreme heterogeneity of neurophysiological experiments impedes the integration, reproduction, interchange, and reuse of diverse neurophysiology data. As other fields of science, such as climate science^3^, astrophysics^4^, and high-energy physics^5^ have demonstrated, community-driven standards for data and metadata are a critical step in creating robust data and analysis ecosystems, as well as enabling collaboration and reuse of data across laboratories. A standardized language for neurophysiology data and metadata (i.e., a data language) is required to enable neuroscientists to effectively describe and communicate about their experiments, and thus share the data.

**Figure 1:**
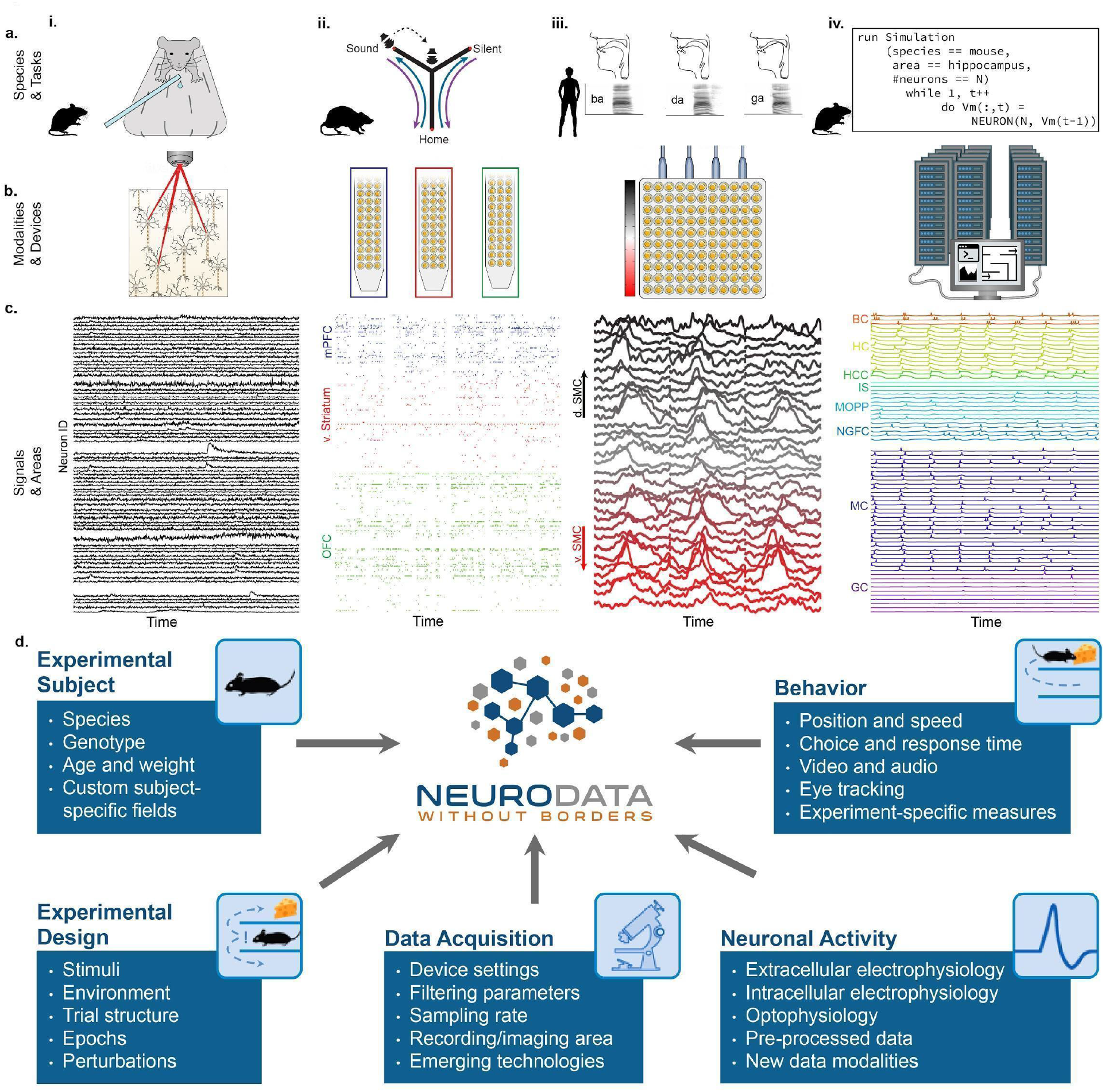
NWB addresses the massive diversity of neurophysiology data and metadata. **a.** Diversity of experimental systems: species and tasks. (**i**). mice performing a visual discrimination task; (**ii**) rats performing a memory-guided navigation task; (**iii**) humans speaking consonant-vowel syllables; (**iv**) biophysically detailed simulations of mouse hippocampus during memory formation. The corresponding acquisition modalities and signals are shown in the corresponding columns in figure **b** and **c**. **b.** Diversity of data modalities and acquisition devices: (**i**) optophysiological Ca^2+^ imaging with 2-photon microscope; (**ii**) intra-cortical extracellular electrophysiological recordings with polytrodes in multiple brain areas (indicated by color, see **cii**); (**iii**) cortical surface electrophysiology recordings with electrocorticography grids; (**iv**) high-performance computing systems for large-scale, biophysically detailed simulations of large neural networks. **c.** Diversity of signals and areas: (**i**) Ca^2+^ signals as a function of time from visually identified individual neurons in primary visual cortex (V1)(Mallory et al., 2021); (**ii**) spike-raster (each tick demarcates the time of an action potential) from simultaneously recorded putative single-units after spike-sorting of extracellular signals from medial prefrontal cortex (mPFC; blue), ventral striatum (v. Striatum, red), and orbital frontal cortex (OFC, green)(color corresponds to **b.ii**)(Kastner et al., 2020); (**iii**) high-gamma band activity from electrodes over the speech sensorimotor cortex (SMC), with dorsal-ventral distance from Sylvian fissure color coded red-to-black (color corresponds to **b.iii**) (Bouchard et al., 2013); (**iv**) simulated intra-cellular membrane potentials from different cell-types from large-scale biophysical simulation of the hippocampus (BC, Basket Cell; HC, Hilar Interneuron (with axon associated with the) Perforant Path; HCC, Hilar Interneuron (with axon associated with the) Commissural/Associational Path; IS, Interneuron-Specific Interneuron; MCPP, medial Perforant Path; NGFC, neurogliaform cell; MC, mossy cell; GC, granule cell](Raikov and Soltesz, unpublished data). **d.** Neurodata Without Borders (NWB) provides a robust, extensible, and maintainable software ecosystem for standardized description, storage, and sharing of the diversity of experimental subjects, behaviors, experimental designs, data acquisition systems, and measures of neural activity exemplified in **a - c**.

The extreme heterogeneity of neurophysiology experiments is exemplified in **Figure 1**. Diverse experiments are designed to investigate a variety of neural functions, including sensation, perception, cognition, and action. Tasks include running on balls or treadmills (e.g., pictures, **Fig. 1i**)^6^, memory-guided navigation of mazes (**Fig. 1ii)**^7^, production of speech (**Fig. 1iii)**^8^, and memory formation (**Fig. 1iv**). The use of different species in neuroscience is driven, in part, by the applicability of specific neurophysiological recording techniques (**Fig. 1b**). For example, the availability of genetically modified mice makes this species ideal to monitor the activity of genetically defined neurons using calcium sensors (e.g., with GCaMP; optophysiology, ‘o-phys’) (**Fig. 1bi,ci**). On the other hand, intracranially implanted electrophysiology probes (‘e-phys’) with large numbers of electrodes enable monitoring the activity of many single neurons at millisecond resolution from different brain regions simultaneously in freely behaving rats (**Fig. 1bii, cii**)^7^. Likewise, in human epilepsy patients, arrays of electrodes on the cortical surface (i.e., electrocorticography, ECoG) provides direct electrical recording of mesoscale neural activity at high-temporal resolution across multiple brain areas (e.g., speech sensorimotor cortex ‘SMC’; **Fig. 1biii, ciii**)^8^. Additionally, to understand the intracellular functioning of single neurons, scientists measure membrane potentials (ic-ephys), e.g., via patch clamp recordings (see **Supplementary Material 1)**. As a final example, to study the detailed workings of complete neural circuits, supercomputers are used for biophysically detailed simulation of the intracellular membrane potentials of a large variety of neurons organized in complex networks^9,10^ (**Fig. 1biv, civ**).

Although the heterogeneity described above is most evident across labs, it is present in a reduced form within single labs; lab members can use new equipment or different techniques in custom experiments to address specific hypotheses. As such, even within the same laboratory, storage and descriptions of data and metadata often vary greatly between experiments, making archival sharing and reuse of data a significant challenge. Across species and tasks, different acquisition technologies measure different neurophysiological quantities from multiple spatial locations over time. Thus, the numerical data itself can commonly be described in the form of space-by-time matrices, the storage of which has been optimized (for space and rapid access) by computer scientists for decades. It is the immense diversity of metadata required to turn those numbers into knowledge that presents the outstanding challenge.

Scientific data must be thought of in the context of the entire data lifecycle, which spans planning, acquisition, processing, and analysis to publication and reuse^11^. In this context, a “data ecosystem” is a shared market for scientific data, software, and services that are able to work together. Such an ecosystem for neurophysiology would empower users to integrate software components and products from across the ecosystem to address complex scientific challenges. Foundational to realizing a data ecosystem is a common ‘language’ that enables seamless exchange of data and information between software components and users. Here, the principles of Findable, Accessible, Interoperable, and Reusable (i.e., FAIR)^12^ data management and stewardship are widely accepted as essential to ensure that data can flow reliably between the components of a data ecosystem. Traditionally, data standards are often understood as rigid and static data models and formats. Such standards are particularly useful to enable the exchange of specific data types (e.g., image data), but are insufficient to address the diversity of data types generated by constantly evolving experiments. Together, these challenges and requirements necessitate a conceptual departure from the traditional notion of a rigid and static data standard. That is, we need a “language” where fundamental structures can be reused and combined in new ways to express novel concepts and experiments. A data language for neurophysiology will enable precise communication about neural data that can co-evolve with the needs of the neuroscience community.

We created the Neurodata Without Borders (NWB) data language (i.e., a standardized language for describing data) for neurophysiology to address the challenges described above. NWB(v2) accommodates the massive heterogeneity and evolution of neurophysiology data and metadata in a unified framework through the development of a novel data language that can co-evolve with neurophysiology experiments. We demonstrate this through the storage of multimodal neurophysiology data, and derived products, in a single NWB file with easy visualization tools. This generality was enabled by the development of a robust, extensible, and sustainable software architecture based on our Hierarchical Data Modeling Framework (HDMF)^13^. To facilitate new experimental paradigms, we developed methods for creating and sharing NWB Extensions that permit the NWB data language to co-evolve with the needs of the community. NWB is foundational for the Distributed Archives for Neurophysiology Data Integration (DANDI) data repository to enable collaborative data sharing and analysis. Together, NWB and DANDI make neurophysiology data FAIR. Indeed, NWB is integrated with a growing ecosystem of state-of-the-art analysis tools to provide a unified storage platform throughout the data life cycle. Through extensive and coordinated efforts in community engagement, software development, and interdisciplinary governance, NWB is now being utilized by more than 53 labs and research organizations. Across these groups, NWB is used for all neurophysiology data modalities collected from species ranging from flies to humans during diverse tasks. Together, the capabilities of NWB provide the basis for a community-based neurophysiology data ecosystem. The processes and principles we utilized to create NWB provide an exemplar for biological data ecosystems more broadly.

## Results

### NWB enables unified description and storage of multimodal data and derived products

NWB files contain all of the measurements for a single experiment, along with all of the necessary metadata to understand that data. Neurophysiology experiments often contain multiple simultaneous streams of data, e.g., via simultaneous recording of neural activity, sensory stimuli, behavioral tracking, and direct neural modulation. Furthermore, neuroscientists are increasingly leveraging multiple neurophysiology recording modalities simultaneously (e.g., ephys and ophys), which offer complementary information not achievable in a single modality. These distinct raw data input types often require processing, further expanding the multiplicity of data types that need to be described and stored.

A key capability of NWB is to describe and store many data sources (including neurophysiological recordings, behavior, and stimulation information) in a unified way that is readily analyzed with all time bases aligned to a common clock. For each data source, raw acquired signals and/or preprocessed data can be stored in the same file. **Figure 2a** illustrates a workflow for storing and processing electrophysiology and optical physiology in NWB^14,15^. Raw voltage traces (**Fig. 2a, top**) from an extracellular electrophysiology recording and image sequences from an optical recording (**Fig. 2a, bottom**) can both be stored in the same NWB file, or separate NWB files synchronized to each other. Extracellular electrophysiology data often goes through spike sorting, which processes the voltage traces into putative single units and action potential (a.k.a., spikes) times for those units (**Fig. 2a, top**). The single unit spike times can then also be written to the NWB file. Similarly, optical physiology is generally processed using segmentation algorithms to identify regions of the image that correspond to neurons and extract fluorescence traces for each neuron (**Fig. 2a, bottom**). The fluorescence traces can also be stored in the NWB file, resulting in raw and processed data for multiple input streams. The timing of these streams is each defined separately, allowing streams with different sampling rates and starting times to be registered to the same common clock. As illustrated in **Figure 1d**, NWB can also store raw and processed behavioral data as well as stimuli, such as animal location and amplitude/frequency of sounds. The multi-modal capability of NWB is critical for capturing the diverse types of data simultaneously acquired in many neurophysiology experiments, particularly if those experiments involve multiple simultaneous neural recording modalities.

**Figure 2.**
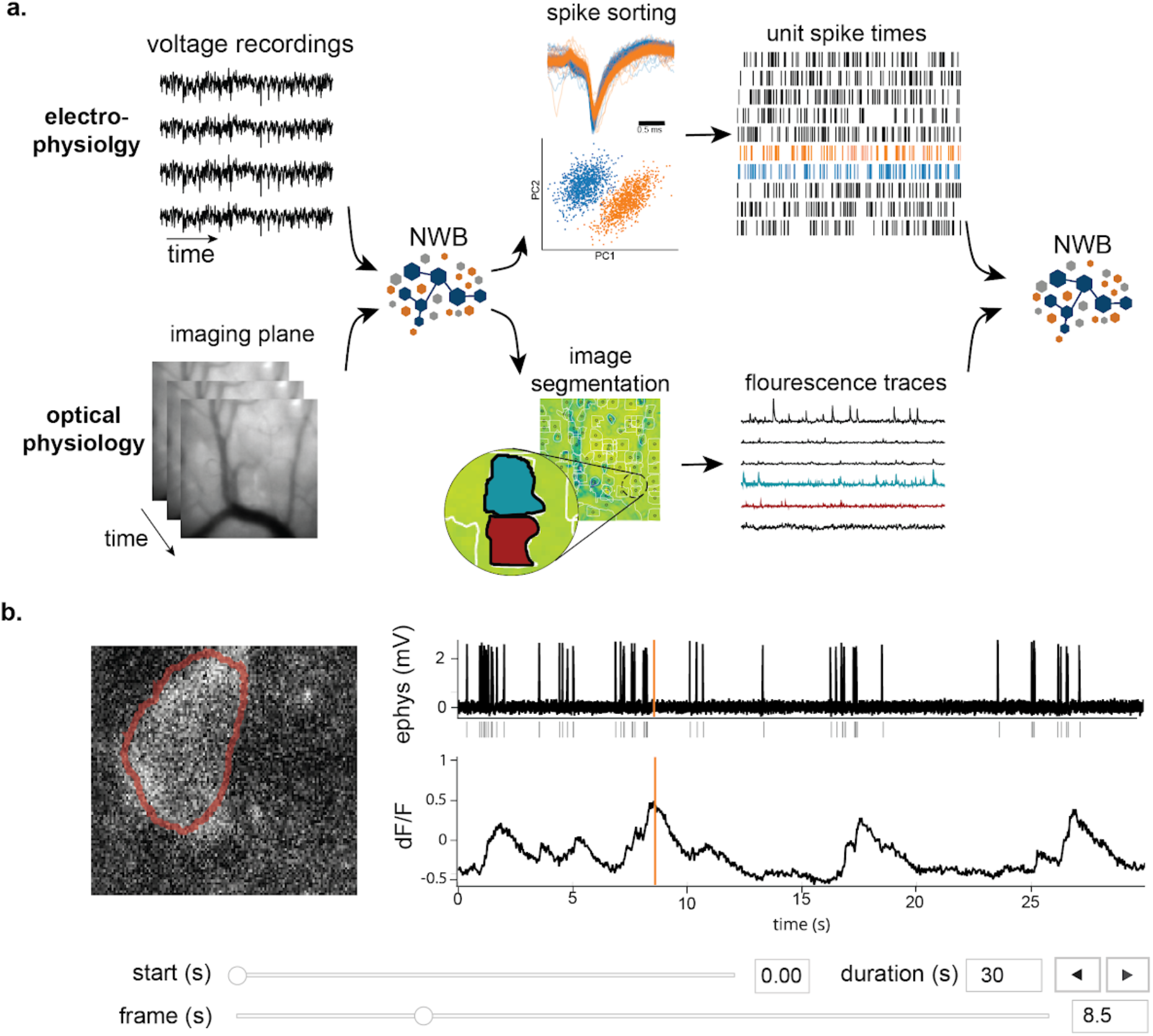
NWB enables unified description and storage of multimodal raw and processed data. **a.** Example pipelines for extracellular electrophysiology and optical physiology demonstrate how NWB facilitates data processing. For extracellular electrophysiology (top), raw acquired data is written to the NWB file. The NWB ecosystem provides interfaces to a variety of spike sorters that extract unit spike times from the raw electrophysiology data. The spike sorting results are then stored into the same NWB file (bottom). Separate experimental data acquired from an optical technique is converted and written to the NWB file. Several modern software tools can then be used to process and segment this data, identifying regions that correspond to individual neurons, and outputting the fluorescence trace of each putative neuron. The fluorescence traces are written to the same NWB file. NWB handles the time alignment of multiple modalities, and can store multiple modalities simultaneously, as shown here. The NWB file also contains essential metadata about the experimental preparation. **b.** NWBWidgets provides visualizations for the data within NWB files with interactive views of the data across temporally aligned data types. Here, we show an example dashboard for simultaneously recorded electrophysiology and imaging data. This interactive dashboard shows on the left the acquired image and the outline of a segmented neuron (red) and on the right a juxtaposition of extracellular electrophysiology, extracted spike times, and simultaneous fluorescence for the segmented region. The orange line on the ephys and dF/F plots indicate the frame that is shown to the left. The controls shown at the bottom allow a user to change the window of view and the frame of reference within that window.

Having pre-synchronized data in the same format enables faster and less error-prone development of analysis and visualizations tools that provide simultaneous views across multiple streams. **Figure 2b** shows an interactive dashboard for exploring a dataset of simultaneously recorded optical physiology and electrophysiology data published by the Allen Institute^14^. This dashboard illustrates the simultaneous exploration of five data elements all stored in a single NWB file. The microscopic image panel (**Fig. 2b, far left)** shows a frame of the video recorded by the microscope. The red outline overlaid on that image shows the region-of-interest where a cell has been identified by the experimenter. The fluorescence trace (dF/F) shows the activation of the region-of-interest over time. This activity is displayed in line with electrophysiology recordings of the same cell (ephys), and extracted spikes (below ephys). Interactive controls **(Fig. 2b, bottom)** allow a user to explore the complex and important relationship between these data sources.

Visualizations of multiple streams of data is a common need across different types of neurophysiology data. Another NWBWidgets dashboard is described in^16^, which demonstrates a dashboard for viewing human body position tracking with simultaneously acquired ECoG data, as well as a panel for viewing the 3D position of electrodes on the participant’s brain. Dashboards for specific experiment types can be constructed using NWBWidgets, a library for interactive web-based visualizations of NWB data, that provides tools for navigating and combining data across modalities.

### The NWB software architecture modularizes and integrates all components of a data language

Neuroscientists use NWB through a core software stack (**Fig. 3b**) with four modularized components: the specification language, the data standard schema, data use APIs, and storage backends (**Fig. 3a**) The identification and modularization of these components was a core conceptual advance of the NWB software. This software architecture provides flexible accommodation of the heterogenous use cases and needs of NWB users. Modularizing the software in this way allows extending the schema to handle new types of data, to implement APIs in new programming languages, and to store NWB using different backends, all while maintaining compliance with NWB and providing a stable interface for users to interact with.

**Figure 3:**
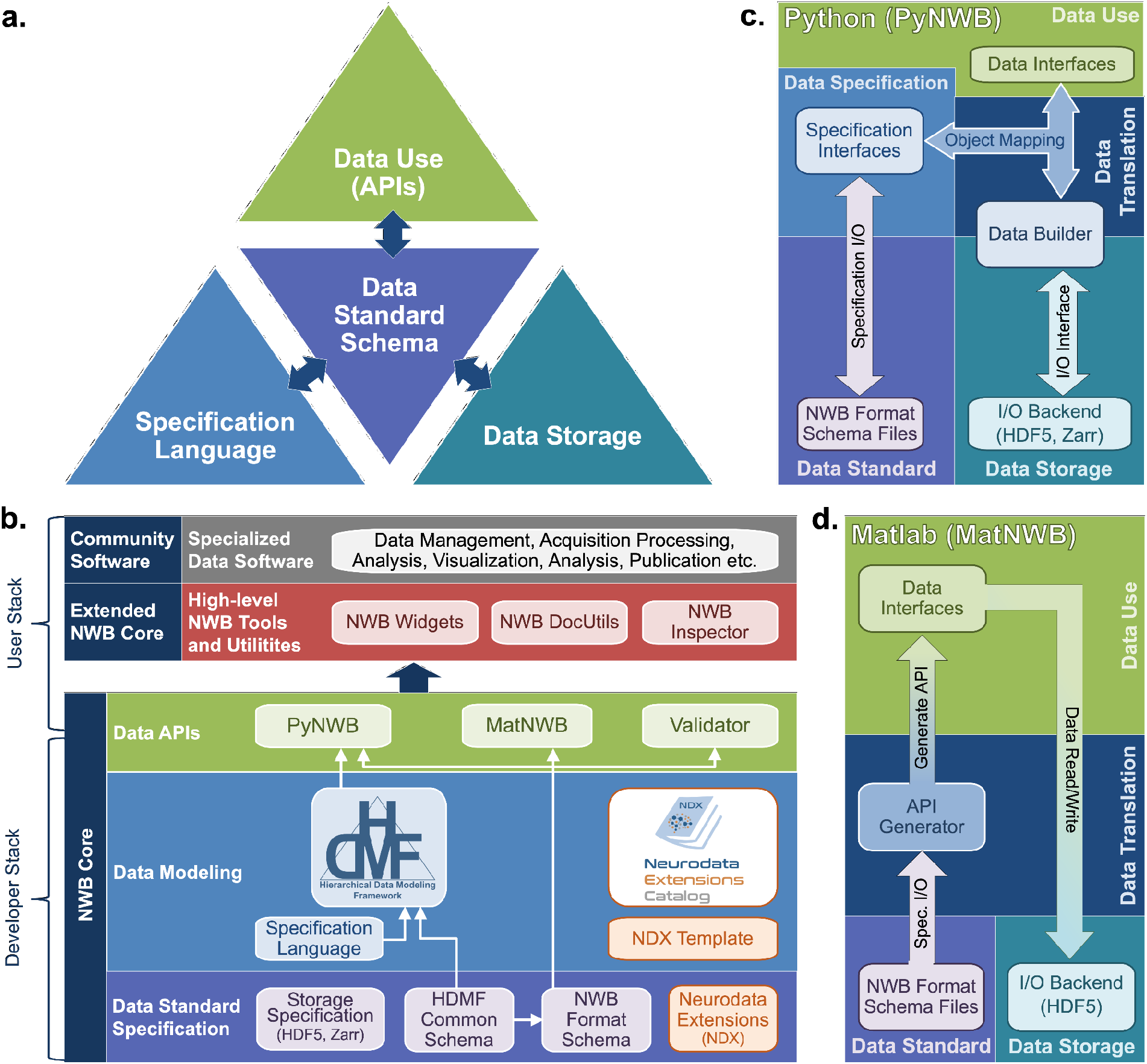
The NWB software architecture modularizes and integrates all components of a data language. **a.** Illustration of the main components of the NWB software stack consisting of: i) the specification language (light blue) to describe data standards, ii) the data standard schema (lilac), which uses the specification language to formally define the data standard, iii) the data storage (blue gray) for translating the data primitives (e.g., groups and datasets) described by the schema to/from disk, and iv) the APIs to enable users to easily read and write data using the standard. Additional data translation components (dark blue arrows) defined in the software then insulate and separate these four main components to enable the individual components to evolve while minimizing impacts on the other components. For example, by insulating the schema from the storage we can extend the standard schema without having to modify the data storage and conversely also integrate new storage backends without having to modify the standard schema. **b.** Software stack for defining and extending the NWB data standard and creating and using NWB data files. The software stack covers all aspects of data standardization: i) data specification, ii) data modeling, iii) data storage, iv) data APIs, v) data translation, and vi) tools. Depending on their role, different stakeholders typically interact with different subsets of the software ecosystem. End users typically interact with the data APIs (green) and higher-level tools (red, gray) while tool developers typically interact with the data APIs and data modeling layers (green, blue). Working groups and developers of extensions then typically interact with the data modeling and data standard specification components. Finally, core NWB developers typically interact with the entire developer stack, from foundational documents (lilac) to data APIs (green). **c.** Software architecture of the PyNWB Python API. PyNWB provides interfaces for interacting with the specification language and schema, data builders, storage backends, and data interfaces. Additional software components (arrows) insulate and formalize the transitions between the various components. The object-mapping-based data translation describes: i) the integration of data interfaces (which describe the data) with the specification (which describes the data model) to generate data builders (which describe the data for storage) and ii) vice versa, the integration of data builders with the specification to create data interfaces. The object mapping insulates the end-users from specifics of the standard specification, builders, and storage, hence, providing stable, easy-to-use interfaces for data use that are agnostic of the data storage and schema. The I/O interface then provides an abstract interface for translating data builders to storage which specific I/O backends must implement. Finally, the specification I/O then describes the translation of schema files to/from storage, insulating the specification interfaces from schema storage details. Most of the data modeling, data translation, and data storage components are general and implemented in HDMF. This approach facilitates the application of the general data modeling capabilities we developed to other science applications and allows PyNWB itself to focus on the definition of data interfaces and functionality that are specific to NWB. **d.** Software architecture of the MatNWB Matlab API. MatNWB generates front-end data interfaces for all NWB types directly from the NWB format schema. This allows MatNWB to easily support updates and extensions to the schema while enabling development of higher-level convenience functions.

First, we describe the specification language used to define hierarchical data models. The YAML-based specification language defines four primitive structures: Groups, Datasets, Attributes, and Links. Each of these primitive structures has characteristics to define their names and parameters (e.g., the allowable shapes of a Dataset). Importantly, these primitives are abstract, and are not tied to any particular data storage backend. The specification language also uses object-oriented principles to define neurodata types that, like classes, can be reused through inheritance and combined through composition to build more complex structures.

The NWB core schema uses the primitives defined in the specification language to define more complicated structures and requirements for particular types of neurophysiology data. For instance, an ElectricalSeries is a neurodata type that defines the data and metadata for an intracranially recorded voltage time series in an extracellular electrophysiology experiment. ElectricalSeries extends the TimeSeries neurodata type, which is a generic structure designed for any measurement that is sampled over time, and defines fields, such as, data, units of measurement, and sample times (specified either with timestamps or sampling rate and start time). ElectricalSeries also requires an electrodes field, which provides a reference to a table of electrodes describing the locations and characteristics of the electrodes used to record the data. The NWB core schema defines many neurodata types in this way, building from generic concepts to specific data elements. The neurodata types have rigorous metadata requirements that ensure a sufficiently rich description of the data for reanalysis. The neurodata types are divided into modules such as ecephys (extracellular electrophysiology), icephys (intracellular electrophysiology), ophys (optical physiology), and behavior. Importantly, the core schema is defined on its own and is agnostic to APIs and programming languages. This allows for the creation of an API in any programming language, which will allow NWB to stay up to date as programming technologies advance.

Application Programming Interfaces (APIs) allow convenient interfaces for writing and reading data according to the NWB schema. The development team maintains APIs in Python (PyNWB) and MATLAB (MatNWB), the two most widely used programming languages in neurophysiology. These APIs (**Fig. 3c,d**) are governed by the NWB schema and use an object-oriented design in which neurodata types (e.g., ElectricalSeries or TowPhotonSeries) are represented by a dedicated interface class. Both APIs are fully compliant with the NWB standard and are, hence, interoperable (i.e., files generated by PyNWB can be read via MatNWB and vice versa). Both APIs also support advanced data Input/Output (I/O) features, such as lazy data read, compression, and iterative data write for data streaming. A key difference in the design of PyNWB and MatNWB is the implementation of the data translation process. PyNWB uses a dynamic data translation process based on data builders (**Fig. 3c**). The data builders are classes that mirror the NWB specification language primitives and provide an interoperability layer where data from different storage backends can be mapped using object mappers into a uniform API. In contrast, MatNWB implements a static translation process that generates the MATLAB API classes automatically from the schema (**Fig. 3d**). The MatNWB approach simplifies updating of the API to support new versions of the NWB schema and extensions, and helps minimize cost for development, but with reduced flexibility in supported storage backend and API. The difference in the data translation process between the APIs (i.e., static vs. dynamic) is a reflection of the different target uses (**Fig. 3c,d**). MatNWB primarily targets data conversion and analysis. In contrast, PyNWB additionally targets integration with data archives and web technologies and is used heavily for development of extensions and exploration of new technologies, such as alternate storage backends and parallel computing libraries.

Finally, the specification of data storage backends deals with translating NWB data models to/from storage on disk. Data storage is governed by formal specifications describing the translation of NWB data primitives (e.g., groups or datasets) to primitives of the particular storage backend format (e.g., HDF5) and is implemented as part of the NWB user APIs. HDF5 is our primary backend, chosen for its broad support across scientific programming languages, its sophisticated tools for handling large datasets, and its ability to express very complex hierarchical structures in relatively few files. The interoperability afforded by the PyNWB builders allows for other backends, and we have a prototype for storing NWB in the Zarr format.

Together, these four components (specification language, standard schema, APIs, and storage backend) and the interaction between them constitute a sophisticated software infrastructure that is applicable beyond neuroscience and could be useful to many other domains. Therefore, we have factored out the domain-agnostic components of each of these four components into a Python software package called the Hierarchical Data Modeling Framework (HDMF) (**Fig. 3b**). Much of the infrastructure described here, including the specification language, fundamental structures of the core schema, base classes for the object mapper and builder layers, and base classes of the PyNWB API are defined in the HDMF package. With its modular architecture and open-source model, the NWB software stack instantiates the NWB data language and makes NWB accessible to users and developers. The NWB software design illustrates the complexity of creating a data language and provides reusable components (e.g., HDMF and the HDMF Common Schema) that can be applied more broadly to facilitate development of data languages for other biological fields in the future.

All NWB software is open source, managed and versioned using Git, and released using a permissive BSD license via GitHub. NWB uses automated continuous integration for testing on all major architectures (MacOS, Windows, and Linux) and all core software can be installed via common package managers (e.g., pip and conda). The suite of NWB core software tools (**Fig. 3b**) enables users to easily read and write NWB files, extend NWB to integrate new data types, and builds the foundation for integration of NWB with community software. NWB data can also be easily accessed in other programming languages (e.g., IGOR or R) using the HDF5 APIs available across modern scientific programming languages.

### NWB enables creation and sharing of extensions to incorporate new use cases

As with all of biology, neurophysiological discovery is driven in large part by new tools that can answer previously unconsidered questions. Thus, a language for neurophysiology data must be able to co-evolve with the experiments being performed and provide customization capability while maintaining stability. NWB enables the creation and sharing of user-defined extensions to the standard that support new and specialized data types (**Fig. 4a1**). Neurodata Extensions (NDX) are defined using the same formal specification language used by the core NWB schema. Extensions can build off of data types defined in the core schema or other extensions through inheritance and composition. This enables the reuse of definitions and associated code, facilitates the integration with existing tools, and makes it easier to contextualize new data types.

**Figure 4:**
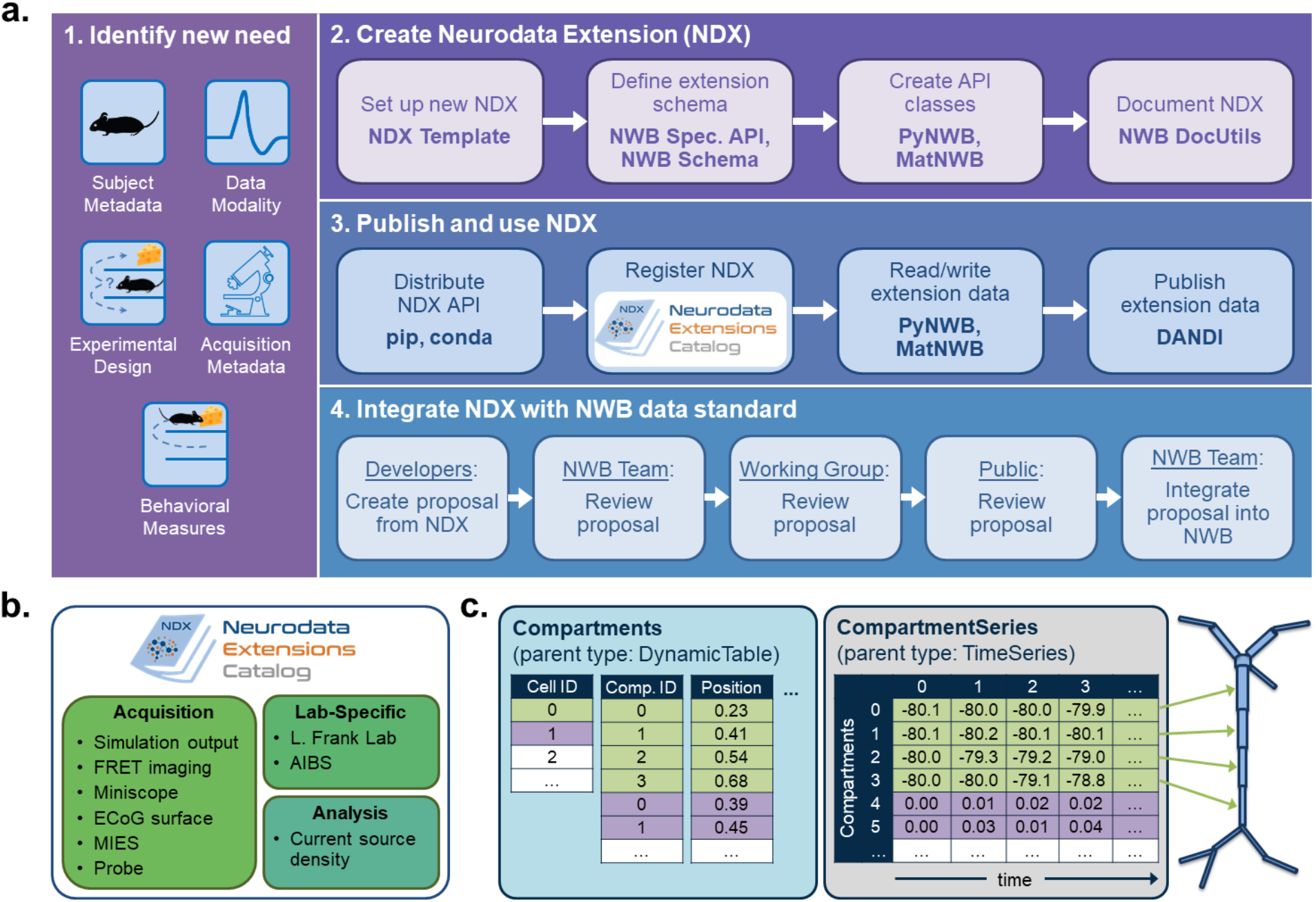
NWB enables creation and sharing of extensions to incorporate new use cases. **a.** Schematic of the process of creating a new neurodata extension (NDX), sharing it, and integrating it with the core NWB data standard. Users first identify the need for a new data type, such as additional subject metadata or data from a new data modality. Users can then use the NDX Template, NWB Specification API, PyNWB/MatNWB data APIs, and NWB DocUtils tools to set up a new NDX, define the extension schema, define and test custom API classes for interacting with extension data, and generate Sphinx-based documentation in common formats, e.g., HTML, PDF. After the NDX is completed, users can publish the NDX on PyPI and conda-forge for distribution via the pip and conda tools, and share extensions via the NDX Catalog, a central, searchable catalog. Users can easily read/write extension data using PyNWB/MatNWB and publish extension data in DANDI and other archives. Finally, extensions are used to facilitate enhancement, maintenance, and governance of the NWB data standard. Users may propose the integration of an extension published in the NDX Catalog with the core standard. The proposal undergoes three phases of review: an initial review by the NWB technology team, an evaluation by a dedicated working group, and an open, public review by the broader community. Once approved, the proposal is integrated with NWB and included in an upcoming version release. **b.** Sampling of extensions currently registered in the NDX catalog. Users can search extensions based on keywords and textual descriptions of extensions. The catalog manages basic metadata about extensions, enabling users to discover and access extensions, comment and make suggestions, contribute to the source code, and collaborate on a proposal for integration into the core standard. While some extensions have broad applicability, others represent data and metadata for a specific lab or experiment. **c.** Example extension for storing simulation output data using the SONATA framework. The new Compartments type extends the base DynamicTable type and contains metadata about each cell and compartment within each cell, such as position and label. The CompartmentSeries type extends the base TimeSeries type and contains a link to the Compartments type to associate each row of its data array with a compartment from the Compartments table.

NWB provides a comprehensive set of tools and services for developing and using neurodata extensions. The NWB Specification API, HDMF DocUtils, and the PyNWB and MatNWB user APIs work with extensions with little adjustment (**Fig. 4a2**). In addition, the NDX Template makes it easy for users to develop new extensions. **Supplementary Material 2** demonstrated the steps outlined in **Fig. 4a2** for the *ndx-simulation-output* extension shown in **Fig. 4c**. The Neurodata Extensions Catalog (**Fig. 4a3**) then provides a centralized listing of extensions for users to publish, share, find, and discuss extensions across the community. **Supplementary Material 3** provides a more detailed overview of the extension workflow as part of the NDX Catalog. Several extensions have been registered in the Neurodata Extensions Catalog (**Fig. 4b**), including extensions to support the storage of the cortical surface mesh of an electrocorticography experiment subject, storage of fluorescence resonance energy transfer (FRET) microscopy data, and metadata from an intracellular electrophysiology acquisition system. The catalog also includes the ndx-simulation-output extension for the storage of the outputs of large-scale simulations. Large-scale network models that are described using the new SONATA format^17^ can be converted to NWB using this extension. The breadth of these extensions demonstrates that NWB will be able to accommodate new experimental paradigms in the future.

As particular extensions gain traction within the community, they may be integrated into the core NWB format for broader use and standardization (**Fig. 4a4**). NWB has a formal, community-driven review process for refining the core format so that NWB can adapt to evolving data needs in neuroscience. The owners of the extension can submit a community proposal for the extension to the NWB Technical Advisory Board, which then evaluates the extension against a set of metrics and best practices published on the catalog website. The extension is then tested and reviewed by both a dedicated working group of potential stakeholders and the general public before it is approved and integrated into the core NWB format. Key advantages of the extension approach are to allow iterative development of extensions and complete implementation and vetting of new data types under several use cases before they become part of the core NWB format. The NWB extension mechanism thus enables NWB to provide a unified data language for all data related to an experiment, allows describing of data from novel experiments, and supports the process of evolving the core NWB standard to fit the needs of the neuroscientific community.

### NWB is foundational for the DANDI data repository to enable collaborative data sharing

Making neurophysiology data accessible supports published findings and allows secondary reuse. To date, many neurophysiology datasets have been deposited into a diverse set of repositories (e.g., CRCNS, Figshare, Open Science Framework, Gin). However, no single data archive provides the neuroscientific community the capacity and the domain specificity to store and access large neurophysiology datasets. Most current repositories have specific limits on data sizes and are often generic, and therefore lack the ability to search using domain specific metadata. Further, for most neuroscientists, these archives often serve as endpoints associated with publishing, while research is typically an ongoing and collaborative process. Few data archives support a collaborative research model that allows data submission prospectively, analysis of data directly in the archive, and opening the conversation to a broader community. Enabling reanalysis of published data was a key challenge identified by the BRAIN Initiative. Together, these issues impede access and reuse of data, ultimately decreasing the return on investment into data collection by both the experimentalist and the funding agencies.

To address these and other challenges associated with neurophysiology data storage and access, we developed DANDI, a Web-based data archive that also serves as a collaboration space for neurophysiology projects (**Fig. 5**). The DANDI data archive (https://dandiarchive.org) is a cloud-based repository for cellular neurophysiology data and uses NWB as its core data language (**Fig. 5b**). Users can organize collections of NWB files (e.g., recorded from multiple sessions) into DANDI datasets (so called Dandisets). Users can view the public Dandisets using a Web browser (**Fig. 5a**) and search for data from different projects, people, species, and modalities. This search is over metadata that has been extracted directly from the NWB files where possible. Users can interact with the data in the archive using a JupyterHub Web interface (**Fig. 5c**) to explore, visualize and analyze data stored in the archive. Using the DANDI Python client, users can organize data locally into the structure required by DANDI as well as download data from and upload data to the archive (**Fig. 5d**). Software developers can access information about Dandisets and all the files it contains using the DANDI Representational State Transfer (REST) API (https://api.dandiarchive.org/). The REST API also allows developers to create software tools and database systems that interact with the archive. Each Dandiest is structured by grouping files belonging to different biosamples, with some relevant metadata stored in the name of each file, and thus aligning itself with the BIDS standard^18^. Metadata in DANDI is stored using the JSON-LD format, thus allowing graph-based queries and exposing DANDI to Google Dataset Search. Dandiset creators can use DANDI as a living repository and continue to add data and analyses to an existing Dandiset. Released versions of Dandisets are immutable and receive a digital object identifier (DOI). The data are presently stored on an Amazon Web Services Public dataset program bucket, enabling open access to the data over the Web, and backed up on institutional repositories. DANDI is working with hardware platforms (e.g., OpenEphys), database software (e.g., DataJoint) and various data producers to generate and distribute NWB datasets to the scientific community.

**Figure 5.**
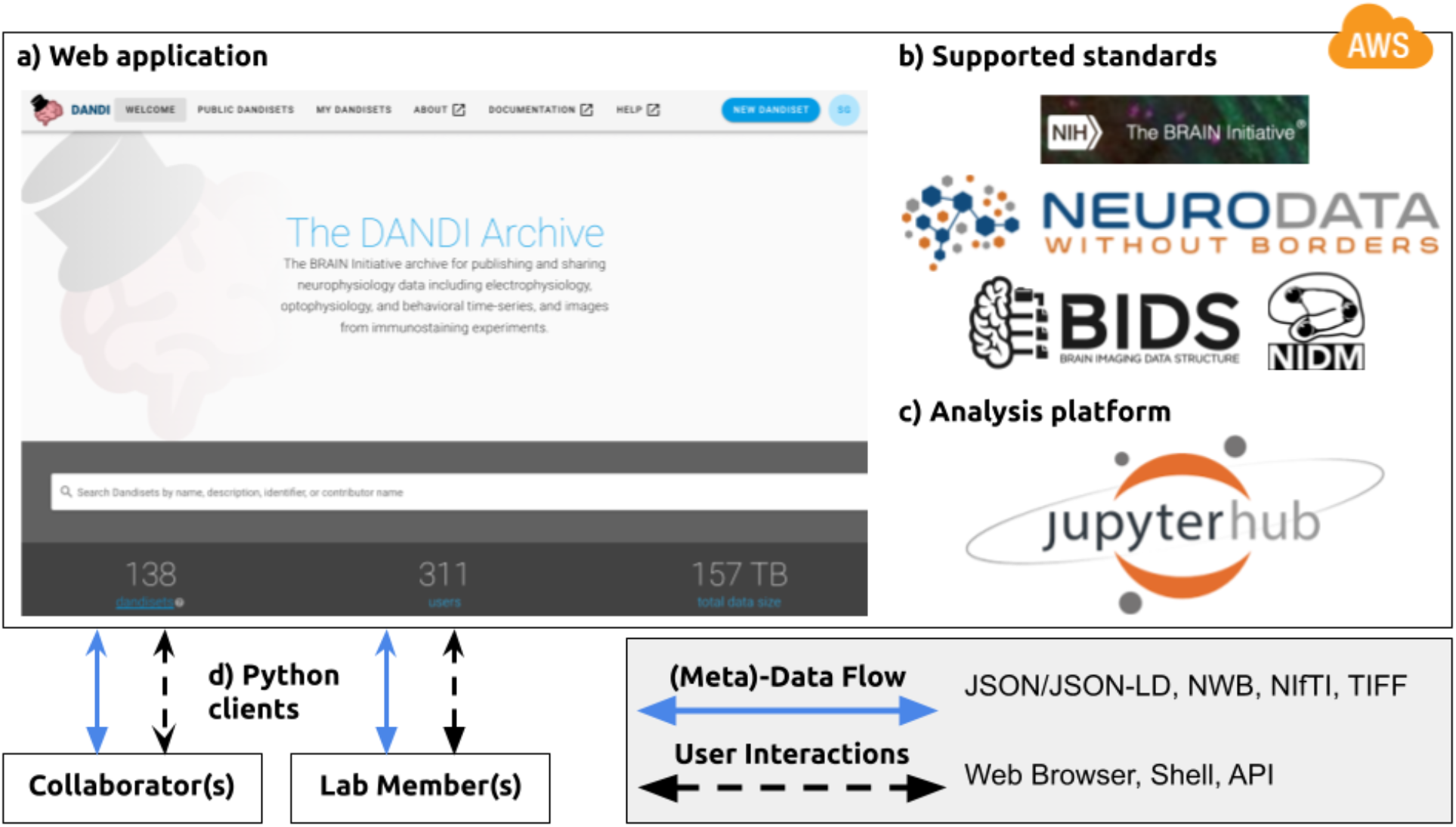
NWB is foundational for the DANDI data repository to enable collaborative data sharing. The DANDI project makes data and software for cellular neurophysiology FAIR. DANDI stores electrical and optical cellular neurophysiology recordings and associated MRI and/or optical imaging data. NWB is foundational for the DANDI data repository to enable collaborative data sharing. **a.** DANDI provides a Web application allowing scientists to share, collaborate, and process data from cellular neurophysiology experiments. The dashboard provides a summary of Dandisets and allows users to view details of each dataset. **b.** DANDI works with US BRAIN Initiative awardees and the neurophysiology community to curate data using community data standards such as NWB, BIDS, and NIDM. DANDI is supported by the US BRAIN Initiative and the Amazon Web Services (AWS) Public Dataset Program. **c.** DANDI provides a JupyterHub interface to visualize the data and interact with the archive directly through a browser, without the need to download any data locally. **d.** Using Python clients and/or a Web browser, researchers can submit and retrieve standardized data and metadata from the archive. The data and metadata use standard formats such as HDF5, JSON, JSON-LD, NWB, NIfTI, and TIFF.

DANDI provides neuroscientists and software developers with a Platform as a Service (PAAS) infrastructure based on the NWB data language and supports interaction via the web browser or through programmatic clients, software, and other services. In addition to serving as a data archive and providing persistence to data, it supports continued evolution of Dandisets. This allows scientists to use the archive to collect data towards common projects across sites, and engage collaborators actively, directly at the onset of data collection rather than at the end. DANDI also provides a computational interface to the archive to accelerate analytics on data and links these Dandisets to eventual publications when generated. The code repositories for the entire infrastructure are available on Github^19^ under an Apache 2.0 license.

### NWB is integrated with state-of-the-art analysis tools throughout the data life cycle

The goal of NWB is to accelerate the rate and improve the quality of scientific discovery through rigorous description and high-performance storage of neurophysiology data. Achieving this goal requires us to consider not just NWB, but the entire data life cycle, including planning, acquisition, processing, and analysis to publication and reuse (**Fig. 6**). NWB provides a common language for neurophysiology data collected using existing and emerging neurophysiology technologies integrated into a vibrant neurophysiology data ecosystem. We describe the software relating to NWB as an “ecosystem,” because it is a marketplace of a diverse set of tools each playing a different role, from data acquisition, to visualization, analysis, and publication. NWB allows scientists to identify an unmet need and contribute new tools to address this need. This is critical to truly make neurophysiology data Findable, Accessible, Interoperable, and Reusable (i.e., FAIR)^12^.

**Figure 6:**
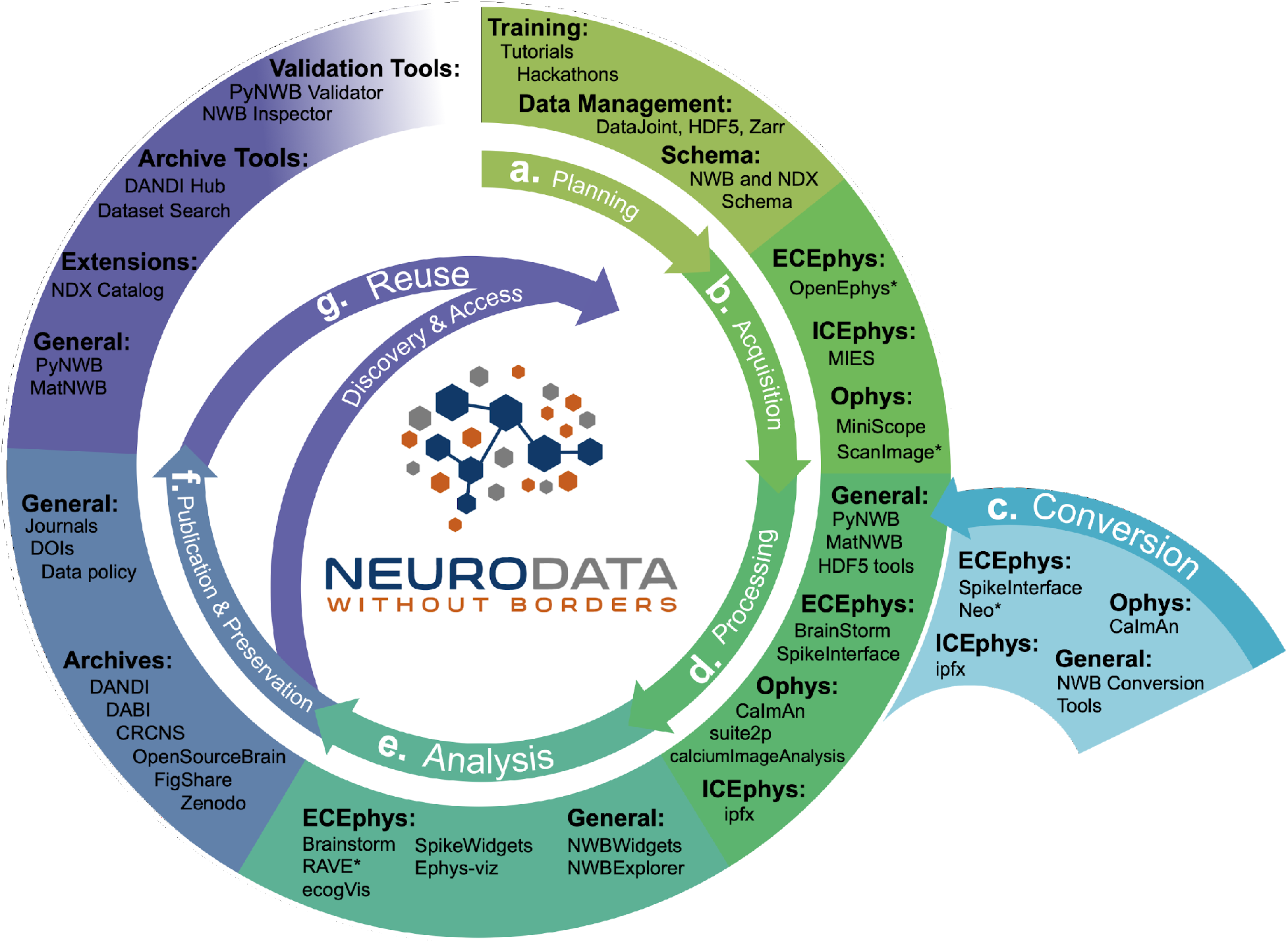
NWB is integrated with state-of-the-art analysis tools throughout the data life cycle. NWB technologies are at the heart of the neurodata lifecycle and applications. Data standards are a critical conduit that facilitate the flow of data throughout the data lifecycle and integration of data and software across all phases (a. to g.) of the data lifecycle. **a.** NWB supports experimental planning through integration with data management, best practices, and by allowing users to clearly define what metadata to collect. **b. c.** NWB supports storage of unprocessed acquired electrical and optical physiology signals, facilitating integration already during data acquisition. NWB is already supported by several acquisition systems (b) as well as a growing set of tools for conversion (c) of existing data to NWB. **d.** Despite its young age, NWB is already supported by a large set of neurophysiology processing software and tools. Being able to access and evaluate multiple processing methods, e.g., different spike sorting algorithms and ROI segmentation methods, is important to enable high-quality data analysis. Through integration with multiple different tools, NWB provides access to broad range of spike sorters, including, MountainSort, KiloSort, WaveClust, and others, and ophys segmentation methods, e.g, CELLMax, CNMF, CNMF-E, and EXTRACT. **e.** For scientific analysis, numerous general tools for exploration and visualization of NWB files (e.g.,. NWBWidgets and NWBExplorer, RRID:SCR_021151) as well as application-specific tools for advanced analytics (e.g., Brainstorm) are accessible to the NWB community. **f. g.** NWB is supported by a growing set of data archives (e.g., DANDI) for publication and preservation of research data. Data archives in conjunction with NWB APIs, validation tools, and the NDX Catalog play a central role in facilitating data reuse and discovery.

NWB supports experiment planning by helping users to clearly define what metadata to collect and how the data will be formatted and managed (**Fig. 6a**). To support data acquisition, NWB allows for the storage of unprocessed acquired electrical and optical physiology signals. Storage of these signals requires either streaming the data directly to the NWB file from the acquisition system or converting data from other formats after acquisition (**Fig. 6b,c**). Some acquisition systems, such as the MIES^20^ intracellular electrophysiology acquisition platform, already support direct recording to NWB and the community is actively working to expand support for direct recording to NWB, e.g., via ScanImage^21^ and OpenEphys^22^. To allow utilization of legacy data and other acquisition systems, a variety of tools exist for converting neurophysiology data to NWB. For extracellular electrophysiology, the SpikeInterface package^23^ provides a uniform API for conversion and processing data that supports conversion for 19+ different proprietary acquisition formats to NWB. For intracellular electrophysiology, the Intrinsic Physiology Feature Extractor (IPFX) package^24^ supports conversion of data acquired with Patchmaster. Direct conversion of raw data to NWB at the beginning of the data lifecycle facilitates data re-processing and re-analysis with up-to-date methods and data re-use more broadly.

The NWB community has been able to grow and integrate with an ecosystem of software tools that offer convenient methods for processing data from NWB files (and other formats) and writing the results into an NWB file (**Fig. 6d**). NWB allows these tools to be easily accessed, compared, and used interoperably. Furthermore, storage of processed data in NWB files allows direct re-analysis of activation traces or spike times via novel analysis methods without having to reproduce time-intensive pre-processing steps. For example, the SpikeInterface API supports export of spike sorting results to NWB across nine different spike sorters and customizable data curation functions for interrogation of results from multiple spike sorters with common metrics. For optical physiology, several popular state-of-the-art software packages, such as CaImAn^25^, suite2p^26^, ciapkg^27^, and EXTRACT ^28^ help users build processing pipelines that segment optical images into regions of interest corresponding to putative neurons, and write these results to NWB.

There is also a range of general and application-specific tools emerging for analysis of neurophysiology data in NWB (**Fig. 6e**). The NWBWidgets^29^ library enables interactive exploration of NWB files via web-based views of the NWB file hierarchy and dynamic plots of neural data, e.g., visualizations of spike trains and optical responses. NWB Explorer^30^ developed by MetaCell in collaboration with OpenSourceBrain, is a web app that allows a user to explore any publicly hosted NWB file and supports custom visualizations and analysis via Jupyter notebooks, as well as use of the NWBWidgets. These tools allow neuroscientists to inspect their own data for quality control, and enable data reusers to quickly understand the contents of a published NWB file. In addition to these general-purpose tools, many application-specific tools, e.g., RAVE^31^, ecogVIS^32^, Brainstorm^333^, Neo^34^ and others are already supporting or are in the process of developing support for analysis of NWB files.

Many journals and funding agencies are beginning to require that data be made FAIR. For publication and preservation (**Fig. 6f**) archives are an essential component of the NWB ecosystem, allowing data producers to document data associated with publications and share that data with others. NWB files can be stored in many popular archives, such as FigShare and Collaborative Research in Computational Neuroscience (CRCNS.org). As described earlier, in the context of the NIH BRAIN Initiative, the DANDI archive has been specifically designed to publish and validate NWB files and leverage their structure for searching across datasets. In addition, several other archives, e.g., DABI^35^ and OpenSourceBrain^36^, are also supporting publication of NWB data.

Data archives also play a crucial role in discovery and reuse of data (**Fig. 6g**). In addition to providing core functionality for data storage and search, archives increasingly also provide compute capacity for reanalysis. For example, DANDI Hub provides users a familiar JupyterHub interface that supports interactive exploration and processing of NWB files stored in the archive. The NWB data APIs, validation, and inspection tools also play a critical role in data reuse by enabling access and ensuring data validity. Finally, the Neurodata Extension Catalog described earlier facilitates accessibility and reuse of data files that use NWB extensions.

NWB integrates with (not competes with) existing and emerging technologies across the data lifecycle, creating a flourishing NWB software ecosystem that enables users to access state-of-the-art analysis tools, share and reuse data, improve efficiency, reduce cost, and accelerate and enable the scientific discovery process. See also **Supplementary Material 4** for an overview of NWB-enabled tools organized by application area and environment. Thus, NWB provides a common language to describe neurophysiology experiments, data, and derived data products that enables users to maintain and exchange data throughout the data lifecycle and access state-of-the-art software tools.

### NWB and DANDI build the foundation for a FAIR neurophysiology data ecosystem

There have been previous efforts to standardize neurophysiology data, such as NWB(v1.0) and NIX^37,38^. While NWB(v1.0) drafted a standard for neurophysiology, it lacked generality which limited its scope, and did not have a reliable and rigorous software strategy and APIs, making it hard to use and unreliable in practice. In contrast, NIX defines a generic data model for storage of annotated scientific datasets, with APIs in C++ and Python and bindings for Java and MATLAB. As such, NIX provides important functionality towards building a FAIR data ecosystem. However, the NIX data model lacks specificity with regard to neurophysiology, leaving it up to the user to define appropriate schema to facilitate FAIR compliance. Due to this lack of specificity, NIX files can also be more varied in structure and naming conventions, which makes it difficult to aggregate across NIX datasets from different labs. In **Table 1** we assess and compare the compliance of different solutions for sharing neurophysiology data with FAIR data principles. The assessment for NIX is based on the INCF review for SPP endorsement ^38^. We also include a more in-depth breakdown of the assessment in **Supplementary Material 5**. With increasing specificity of data models and standard schema––i.e., as we move from general, self-describing formats (e.g., Zarr or HDF5) to generic data models (e.g., NIX) to application-specific standards (e.g., NWB) –– compliance with FAIR principles and rigidness of the data specification increases. In practice, the various approaches often focus on different data challenges. As such, this is not an assessment of the quality of a product per-se, but an assessment of the out-of-the-box compliance with FAIR principles in the context of neurophysiology. For example, while self-describing data formats (like HDF5 or Zarr) lack specifics about (meta)data related to neurophysiology, they provide important technical solutions towards enabling high-performance data management and storage.

**Table 1.**
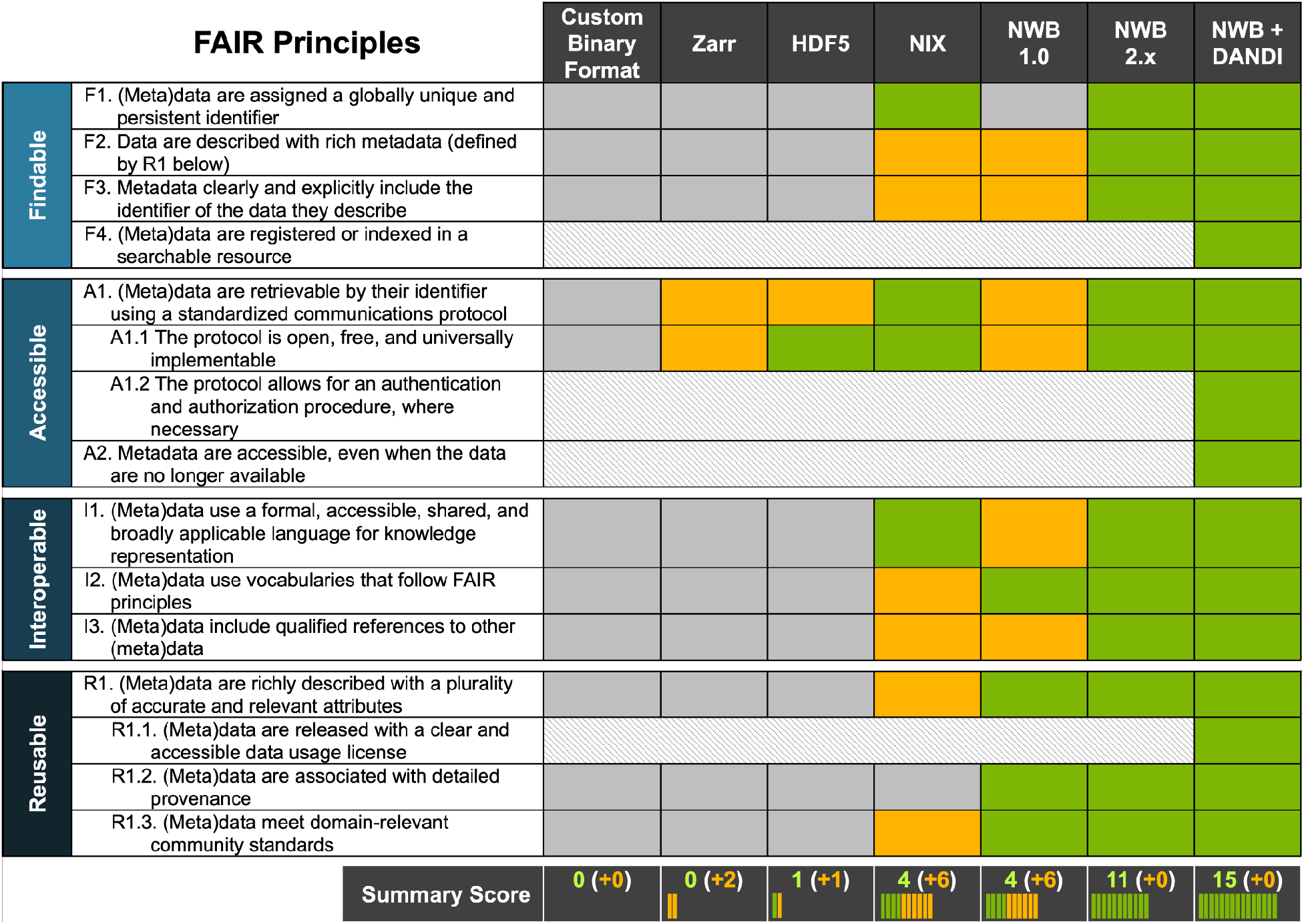
NWB together with DANDI provides an accessible approach for FAIR sharing of neurophysiology data. The table above assesses various approaches for sharing neurophysiology data with regard to their compliance with FAIR data principles. Here cells shown in gray/green indicate non-compliance and compliance, respectively. Cells shown in yellow indicate partial compliance, either due to incomplete implementation or optional support, leaving achieving compliance ultimately to the end user. The larger, shaded blocks indicate areas that are typically not covered by data standards directly but are the role of other resources in a FAIR data ecosystem, e.g., the DANDI data archive.

Complementary to standardization of data, software packages, e.g., Neo^34^, SpikeInterface^23^ and others, aim to simplify programmatic interaction with neurophysiology data in diverse formats and/or tools with diverse programming interfaces (e.g., for spike sorting) by providing common software interfaces for interacting with the data/tools. This strategy provides an important conduit to enable access to and facilitate integration with a diversity of data and tools. However, this approach does not address (nor does it aim to address) the issue of compliance of data with FAIR principles, but it rather aims to improve interoperability between and interaction with a diversity of tools and data formats. Ultimately, standardization of data and creation of common software interfaces are not competing strategies, but are synergistic approaches that together help create a more integrated data ecosystem. Indeed, tools such as SpikeInterface^23^, are an important component of the larger NWB software ecosystem that help create an accessible neurophysiology data ecosystem by making it easier for users to integrate their data and tools with NWB and facilitating access and interoperability of diverse tools.

Data standards build the foundation for an overall data strategy to ensure compliance with FAIR data principles. Ultimately, however, ensuring FAIR data sharing and use depends on an ecosystem of data standards and data management, analysis, visualization, and archive tools as well as laws, regulations, and data governance structures––e.g., the NIH BRAIN Initiative Data Sharing Policy^39^ or the OMB Open Data Policy^40^––all working together^41^. As **Table 1** illustrates, it is ultimately the combination of NWB and DANDI working together that enable compliance with FAIR principles. Here, certain aspects, such as usage licenses (R.1.1), indexing and search (F.4), authenticated access (A1.2), and long-term availability of metadata (A2) are explicitly the role of the archive. As this table shows, together, NWB and DANDI make neurophysiology data FAIR.

### Coordinated Community Engagement, Governance, and Development of NWB

The neurophysiology community consists of a large diversity of stakeholders with vested interests and broad use cases. Inclusive engagement and outreach with the community are central to achieve acceptance and adoption of NWB and to ensure that NWB meets user needs. Thus, development of scientific data languages is as much a sociological challenge as it is a technological challenge. To address this challenge, NWB has adopted a modern open-source software strategy (**Fig. 7a**) with community resources and governance, and a variety of engagement activities.

**Figure 7.**
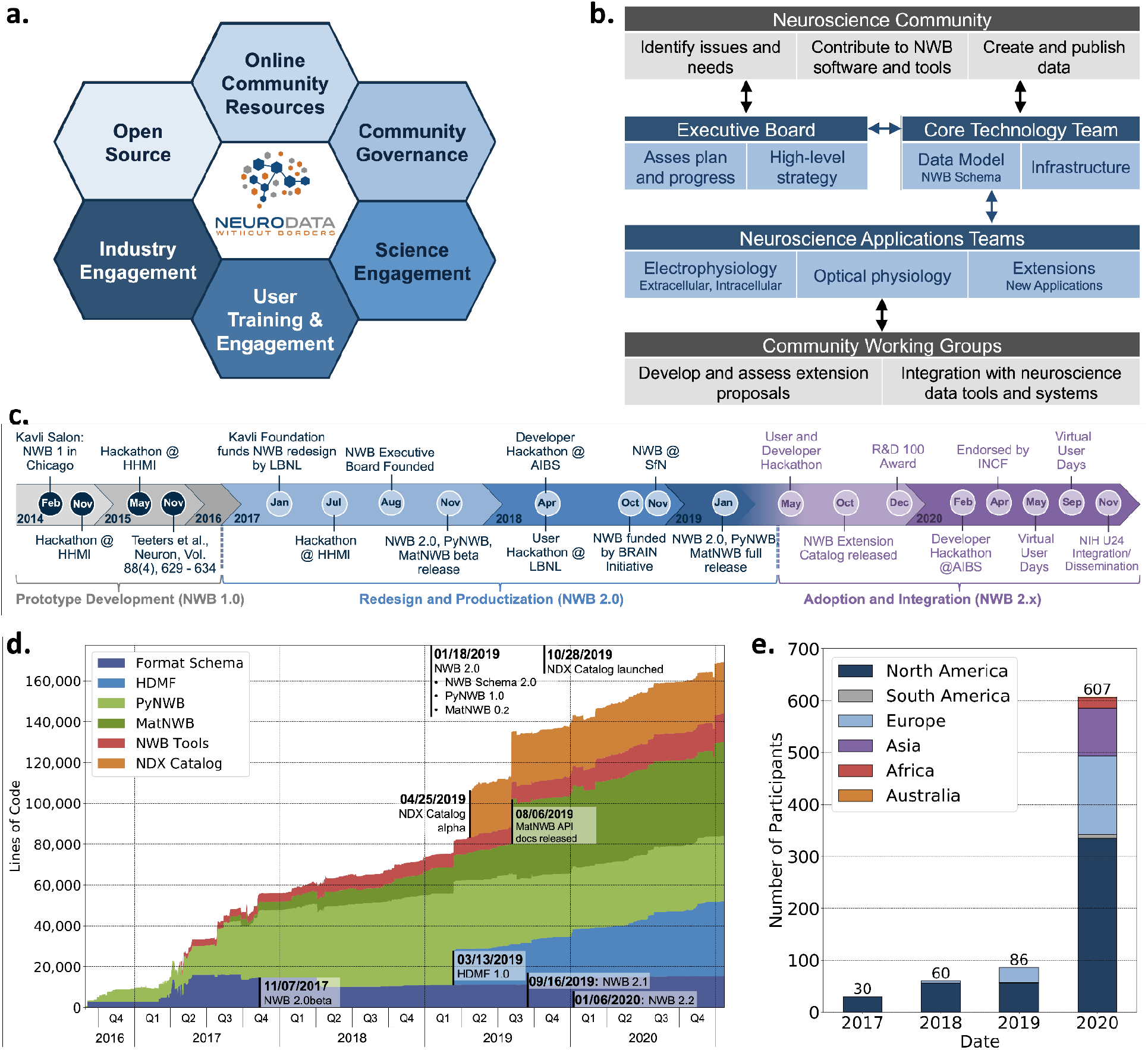
Coordinated Community Engagement, Governance, and Development of NWB. **a.** NWB is open source with all software and documents available online via GitHub and the nwb.org website. NWB provides a broad range of online community resources, e.g., Slack, online Help Desk, GitHub, mailing list, or Twitter, to facilitate interaction with the community and provides a broad set of online documentation and tutorials. NWB uses an open governance model that is transparent to the community. Broad engagements with industry partners (e.g., DataJoint, Kitware, MathWorks, MBFBioscience, Vidrio, CatalystNeuro, etc.) and targeted science engagements with neuroscience labs and tool developers help sustain and grow the NWB ecosystem. Broad user training and engagement activities, e.g., via hackathons, virtual training, tutorials at conferences, or online training resources, aim at facilitating adoption and growing the NWB community knowledge base. **b.** Organizational structure of NWB showing the main bodies of the NWB team (blue boxes) and the community (gray boxes), their roles (light blue/gray boxes), and typical interactions (arrows). **c.** The timeline of the NWB project to date can be roughly divided into three main phases. The initial NWB pilot project (2014-2015) resulted in the creation of the first NWB 1.0 prototype data standard. The NWB 2.0 effort then focused on facilitating use and long-term sustainability by redesigning and productizing the data standard and developing a sustainable software strategy and governance structure for NWB (2017-2019). The release of NWB 2.0 in Jan. 2019 marked the beginning of the third main phase of the project, focused on adoption and integration of NWB with neuroscience tools and labs, maintenance, and continued evolution and refinement of the data standard. **d.** Overview of the growth of core NWB 2.x software in lines of code over time. **e.** Number of participants at NWB outreach and training events over time. In the count we considered only the NWB hackathons and User Days (see **c**.), the 2019 and 2020 NWB tutorial at Cosyne, and 2019 training at the OpenSourceBrain workshop (i.e., not including attendees at presentations at conferences, e.g., SfN).

Execution of this strategy requires coordinating efforts across stakeholders, use cases, and standard technologies to prioritize software development and resolve potential conflicts. Such coordination necessitates a governance structure reflecting the values of the project and the diverse composition of the community (**Fig. 7b**). The NWB Executive Board consists of diverse experimental and computational neuroscientists from the community and serves as the steering committee for developing the long-term vision and strategy. The NWB Executive Board (see Acknowledgements) was established in 2007 as an independent body from the technical team and PIs of NWB grants. The NWB Core Technology Team leads and coordinates the development of the NWB data language and software infrastructure to ensure quality, stability, and consistency of NWB technologies, as well as timely response to user issues. The Core Technology Team reports regularly to and coordinates with the Executive Board. Neuroscience Application Teams, consisting of expert users and core developers lead, engagement with targeted neuroscience areas in electrophysiology, optical physiology, and other emerging applications. These application teams are responsible for developing extensions to the data standard, new features, and technology integration together with Community Working Groups. The working groups allow for an agile, community-driven development and evaluation of standard extensions and technologies, and allow users to directly engage with the evolution of NWB. The broader neuroscience community further contributes to NWB via issue tickets, contributions to NWB software and documentation, and by creating and publishing data in NWB. This governance and development structure emphasizes a balance between the stability of NWB technologies that ensure reliable production software, direct engagement with the community to ensure that NWB meets diverse stakeholder needs, and agile response to issues and emerging technologies.

The balance between stability, diversity, and agility is also reflected in the overall timeline of the NWB project (**Fig. 7c**). The NWB 1.0 prototype focused on evaluation of existing technologies and community needs and development of a draft data standard. Building on this prototype, the NWB 2.0 project initially focused on the redesign and productization of NWB, emphasizing the creation of a sustainable software architecture, reliable data standard, and software ready for use. Following the first full release of NWB 2.0 in January 2019, the emphasis then shifted to adoption and integration of NWB. The goal has been to grow a community and software ecosystem as well as maintenance and continued refinement of NWB.

Together, these technical and community engagement efforts have resulted in a vibrant and growing ecosystem of public NWB data (**Supplementary Material 6**) and tools utilizing NWB. The core NWB software stack has continued to grow steadily since the release of NWB 2.0 in January 2019, illustrating the need for continued development and maintenance of NWB (**Fig. 7d**). See also **Supplementary Martial 7** for an overview of the software release process and history for the NWB schema and APIs. In 2020 more than 600 scientists participated in NWB developer and user workshops and we have seen steady growth in attendance at NWB events over time (**Fig. 7e**). At the same time, the global reach of NWB has also been increasing over time (**Fig. 7e**). The NWB team also provides extensive online training resources, including video and code tutorials, detailed documentation, as well as guidelines and best practices (see **Supplement Material 8, 9** and Methods). Community liaisons provide expert consultation for labs adopting NWB and for creating customized data conversion software for individual labs. As the table in **Supplementary Material 6** shows, despite its young age relative to the neurophysiology community, NWB 2.0 is being adopted by a growing number of neuroscience laboratories and projects led by diverse principal investigators, creating a representative community where users can exchange and reuse data, with NWB as a common data language^42^.

## Discussion

Investigating the myriad functions of the brain across species necessitates a massive diversity and complexity of neurophysiology experiments. This diversity presents an outstanding barrier in meaningful sharing and collaborative analysis of the collected data, and ultimately prevents the data from being FAIR. To overcome this barrier, we developed a data ecosystem based on the Neurodata Without Borders (NWB) data language and software. NWB is being utilized by more than 36 labs to enable unified storage and description of intracellular, extracellular, LFP, ECoG, and Ca^2+^ data in fly, mice, rats, monkeys, humans, and simulations. To support the entire data lifecycle, NWB natively operates with processing, analysis, visualization, and data management tools, as exemplified by the ability to store both raw and pre-processed simultaneous electrophysiology and optophysiology data. Formal extension mechanisms enable NWB to co-evolve with the needs of the community. NWB enables DANDI to provide a data archive that also serves as a collaboration space for neurophysiology projects. Together, these technologies greatly enhance the FAIRness of neurophysiology data.

We argue that there are several key challenges that, until NWB, have not been successfully addressed and which ultimately hindered wide-spread adoption of a common standard by the diverse neurophysiology community. Conceptually, the complexity of the problem necessitates an interdisciplinary approach of neuroscientists, data and computer scientists, and scientific software engineers to identify and disentangle the components of the solution. Technologically, the software infrastructure instantiating the standard must integrate the separable components of user-facing interfaces (i.e., Application Programming Interfaces, APIs), data modeling, standard specification, data translation, and storage format. This must be done while maintaining sustainability, reliability, stability, and ease of use for the neurophysiologist. Furthermore, because science is advanced by both development of new acquisition techniques and experimental designs, mechanisms for extending the standard to unforeseen data and metadata are essential. Sociologically, the neuroscientific community must accept and adopt the standard, requiring coordinated community engagement, software development, and governance. NWB directly addresses these challenges.

### NWB as the *lingua franca* of neurophysiology data

Making neurophysiology data FAIR requires a paradigm shift in how we conceptualize the solution. Scientists need more than a rigid data format, but instead require a flexible data language. Such a language should enable scientists to communicate via data. Natural languages evolve with the concepts of the societies that use them, while still providing a stable basis that enables communication of common concepts. Similarly, a scientific data language should evolve with the scientific research community, and at the same time provide a standardized core that expresses common and established methods and data types. NWB is such a data language for neurophysiology experiments.

There are many parallels between NWB and natural languages as used today. The NWB specification language provides the basic tools and rules for creating the core concepts required to describe neurophysiology data, much like an alphabet and phonetic rules in natural language describe the creation of words. Likewise, the format schema provides the words and phrases (neurodata_types) of the data language and rules for how to compose them to form data documents (NWB files), much like a dictionary and grammatical rules for sentence and document structure. Similarly, flexible data storage methods allow NWB to manage and share data in different forms depending on the application, much like we store natural languages in many different mediums (e.g., via printed books, electronic records, or handwritten notes). User APIs (here HDMF, PyNWB, and MatNWB) provide the community with tools to create, read, and modify data documents and interact with core aspects of the language, similarly to text editors for natural language. NWB Extensions provide a mechanism to create, publish, and eventually integrate new modules into NWB to ensure it co-evolves with the tools and needs of the neurophysiology community, just as communities create new words to communicate emerging concepts. Finally, DANDI provides a cloud-based platform for archiving, sharing, and collaborative analysis of NWB data, much like a bookstore or Wikipedia. Together, these interacting components provide the basis of a data language and exchange medium neuroscience community that enables reproduction, interchange, and reuse of diverse neurophysiology data.

### NWB is community driven, professionally developed, and democratizes neurophysiology

Today, there are many data formats and tools used by the neuroscience community that are not interoperable. Often, formats and tools are specific to the lab and even the researcher. This level of specificity is a major impediment to sharing data and reproducing results, even within the same lab. More broadly, the resulting fragmentation of the data space reinforces siloed research, and makes it difficult for datasets or software to be impactful on a community level. Our goal is for the NWB data language to be foundational in deepening collaborations between the community of neuroscientists. The current NWB software is the result of an intense, community-led, years-long collaboration between experimental and computational neuroscientists with data scientists and computer scientists. Core to the principles and success of NWB is to account for diverse perspectives and use cases in the development process, integration with community tools, and engage in community outreach and feedback collection. NWB is governed by a diverse group to ensure both the integrity of the software and that NWB continues to meet the needs of the neuroscience community.

As with all sophisticated scientific instruments, there is some training required to get a lab’s data into NWB. Several training and outreach activities provide opportunities for the neuroscience community to learn how to most effectively utilize NWB. Tutorials, hackathons and user training events allow us to bring together neuroscientists who are passionate about open data and data management. These users bring their own data to convert or their own tools to integrate, which in turn makes the NWB community more diverse and representative of the overall neuroscience community. NWBs digital presence has accelerated during the COVID pandemic, and has allowed the community to grow internationally and at an exponential rate. Updates on Twitter and the website (www.nwb.org), tutorials on YouTube, and free virtual hackathons, all are universally accessible and have helped achieve a global reach, interacting with scientists from countries that are too often left out. Together, these outreach activities combined with NWB and DANDI democratizes both neurophysiology data and analysis tools, as well as the extracted insights.

### The Future of NWB

To address the next frontier in grand challenges associated with understanding the brain, the neuroscience community must integrate information across experiments spanning several orders of magnitude in spatial and temporal scales^43–45^. This issue is particularly relevant in the current age of massive neuroscientific data sets generated by emerging technologies from the US BRAIN Initiative, Human Brain Project, and other brain research initiatives worldwide. Advanced data processing, machine learning, and artificial intelligence algorithms are required to make discoveries based on such massive volumes of data^43,44,46^. Currently, different domains of neuroscience (e.g., genomic/transcriptomic, anatomy, neurophysiology, etc.,) are supported by standards that are not coordinated. Building bridges across neuroscience domains will necessitate interaction between the standards, and will require substantial future efforts. There are nascent activities for compatibility between NWB and the Brain Imaging Data Structure (BIDS), e.g., as part of the BIDS human intracranial neurophysiology ECoG/iEEG extension^47^, but further efforts in this and other areas are needed.

It is notoriously challenging to make neurophysiology data FAIR. Together, the NWB data language and the NWB-based DANDI data archive support a data science ecosystem for neurophysiology. NWB provides the underlying cohesion of this ecosystem through a common language for the description of data and experiments. However, like all languages, NWB must continue to adapt to accommodate advances in neuroscience technologies and the evolving community using that language. As adoption of NWB continues to grow, new needs and opportunities for further harmonization of metadata arise. A key ongoing focus area is on development and integration of ontologies with NWB to enhance specificity, accuracy, and interpretability of data fields. For example, there are NWB working groups on genotype and spatial coordinate representation, as well as the INCF Electrophysiology Stimulation Ontology working group^48^. Another key area is extending NWB to new areas, such as the ongoing working groups on integration of behavioral task descriptions with NWB (e.g., based on BEADL^49^) and enhanced integration of simulations with NWB. We strongly advocate for funding support of all aspects of the data-software life cycle (development, maintenance, integration, and distribution) to ensure the neuroscience community fully reaps the benefits of investment into neurophysiology tools and data acquisition.

### Core design principles and technologies for biological data languages

The problems addressed by NWB technologies are not unique to neurophysiology data. Indeed, as was recently discussed in Powell 2021^50^, lack of standards in genomics data is threatening the promise of that data. Many of the tools and concepts of the NWB data language can be applied to enhance standardization and exchange of data in biology more broadly. For example, the specification language, HDMF, the concept of extensions and the extension catalog are all general and broadly applicable technologies. Therefore, the impact of the methods and concepts we have described here has the potential to extend well beyond the boundaries of neurophysiology.

We developed design and implementation principles to create a robust, extensible, maintainable, and usable data ecosystem that embraces and enables FAIR data science across the breadth of neurophysiology data. Across biology, experimental diversity and data heterogeneity are the rule, not the exception^2^. Indeed, as biology faces the daunting frontier of understanding life from atoms to organisms, the complexity of experiments and multimodality of data will only increase. Therefore, the principles developed and deployed by NWB may provide a blueprint for creating data ecosystems across other fields of biology.

## Methods

### NWB GitHub Organizations

All NWB software is available open source via the following three GitHub organizations. The Neurodata Without Borders^51^ GitHub organization is used to manage all software resources related to core NWB software developed by the NWB developer community, e.g., the PyNWB and MatNWB reference APIs. The HDMF development^52^ Github organization is used to publish all software related to the Hierarchical Data Modeling Framework (HDMF), including, HDMF, HDMF DocUtils, HDMF Common Schema and others. Finally, the NWB Extensions^53^ GitHub organization is used to manage all software related to the NDX Catalog, including all extension registrations. Note, the catalog itself only stores metadata about NDXs, the source code of NDXs are often managed by the creators in dedicated repositories in their own organizations.

### HDMF Software

HDMF software is available on GitHub using an open BSD licence model.

**<u>H</u>ierarchical <u>D</u>ata <u>M</u>odeling <u>F</u>ramework (HDMF)** is a Python package for working with hierarchical data. It provides APIs for specifying data models, reading and writing data to different storage backends, and representing data with Python objects. HDMF builds the foundation for the implementation of PyNWB and specification language. **[Source]**^54^ **[Documentation]**^55^ **[Web]**^56^

**HDMF Documentation Utilities (hdmf-docutils)** are a collection of utility tools for creating documentation for data standard schema defined using HDMF. The utilities support generation of reStructuredText (RST) documents directly from standard schema which can be compiled to a large variety of common document formats (e.g., HTML, PDF, epub, man and others) using Sphinx. **[Source]**^57^

**HDMF Common Schema** defines a collection of common reusable data structures that build the foundation for modeling of advanced data formats, e.g., NWB. APIs for the HDMF common data types are implemented as part of the `hdmf.common` module in the HDMF library. **[Source]**^58^ **[Documentation]**^59^

**HDMF Schema Language** provides an easy-to-use language for defining hierarchical data standards. APIs for creating and interacting with HDMF schema are implemented in HDMF. **[Documentation]**^60^

### NWB Software

NWB software is available on GitHub using an open BSD licence model.

**PyNWB** is the Python reference API for NWB and provides a high-level interface for efficiently working with Neurodata stored in the NWB format. PyNWB is used by users to create and interact with NWB and neuroscience tools to integrate with NWB. **[Source]**^61^ **[Documentation]**^62^

**MatNWB** is the MATLAB® reference API for NWB and provides an interface for efficiently working with Neurodata stored in the NWB format. MatNWB is used by both users and developers to create and interact with NWB and neuroscience tools to integrate with NWB. **[Source]**^63^ **[Documentation]**^64^

**NWBWidgets** is an extensible library of widgets for visualizing NWB data in a Jupyter notebook (or lab). The widgets support navigation of the NWB file hierarchy and visualization of specific NWB data elements. **[Source]**^65^

**NWB Schema** defines the complete NWB data standard specification. The schema is a collection of YAML files in the NWB specification language describing all `neurodata_types` supported by NWB and their organization in an NWB file. **[Source]**^66^ **[Documentation]**^67^

**NWB Schema Language** is a specialized variant of the HDMF schema language. The language includes minor modifications (e.g., use of the term `neurodata_type` instead of `data_type`) to make the language more intuitive for neuroscience users, but it is otherwise identical to the HDMF schema language. Dedicated interfaces for creating and interacting with NWB schema are available in PyNWB. **[Documentation]**^68^

**NWB Storage** defines the mapping of NWB specification language primitives to HDF5 for storage of NWB files. **[Documentation]**^69^

**Neurodata Extensions Catalog (NDX Catalog)** is a community-led catalog of Neurodata Extensions (NDX) to the NWB data standard. All extensions mentioned in the text can be accessed directly via the catalog. **[Source]**^53^ **[Online]**^70^

**NWB Extensions Template (ndx-template)** provides an easy-to-use template based on the Cookiecutter library for creating Neurodate Extensions (NDX) for the NWB data standard. **[Source]**^71^

**NWB Staged Extensions** is a repository for submitting new extensions to the NDX catalog [Source]^72^

### DANDI

The DANDI archive was created by developing and integrating several opensource projects and BRAIN Initiative data standards (NWB, BIDS, NIDM). The Web browser application is built using the VueJS framework and the DANDI command line interface is built using Python and PyNWB. The initial backend of the archive was built on top of the Girder data management system, and is transitioning to a Django-based framework. The DANDI analysis hub is built using Jupyterhub deployed over a Kubernetes cluster. The different components of the archive are hosted on Amazon Web Services and the Heroku platform. The code repositories for the entire infrastructure are available on Github^19^ under an Apache 2.0 license.

## Acknowledgements

Research reported in this publication was supported by the following grants and institutions. OR, AT, RL, and KEB were supported by the National Institute Of Mental Health of the National Institutes of Health under Award Number R24MH116922 (PI: O. Ruebel) as well as through additional support by the Kavli Foundation (PI: KEB). BD and IS were supported by the BRAIN Initiative under award number U19-NS104590 to IS and by the Kavli Foundation. SG and BD were supported by the NIH BRAIN Initiative under award number 1R24MH117295-01A1. KS is supported by HHMI. Additional support for NWB was also provided by the Simons Foundation for the Global Brain grant 521921 to LF, and with additional funding from the Kavli Foundation and Simons Foundation for MatNWB to KS. KEB was additionally supported by the Weill Neurohub. CatalystNeuro is also supported by the Simons Foundation.

We thank the diverse participants of the NWB hackathons and the global NWB user and developer community for their feedback and contributions. We thank Michael Grauer, Matt McCormick, Jean-Christophe Fillion-Robin, Doruk Ozturk, Roni Choudhury and Pamela Baker for early work on CI/CD. We thank the current and former members of the NWB Executive Board for their guidance and support: Kristofer Bouchard, Bing Brunton, Elizabeth Buffalo, Anne Churchland, Loren Frank, Satrajit Ghosh, Adam Kepecs, Lydia Ng, Huib Mansvelder, Ueli Rutishauser, Karel Svoboda, Christof Koch, Friedrich Sommer, Markus Meister, and Katrin Amunts.

## Author Contributions

Conceptualization: KEB, OR

Methodology: OR, AT, RL, SG, BKD

Software: RL, AT, OR, LN Resource: KEB, OR, IS, LF, KS

Writing - Original Draft: KEB, BKD, SG, RL, OR, AT

Writing - Review & Editing: KEB, BKD, SG, RL, OR, AT, LF, IS, KS

Visualization: KEB, BKD, SG, RL, OR, AT

Supervision: KEB, OR

Project Administration: KEB, OR

Funding Acquisition: KEB, OR, SG, IS, LF, KS

