## Supplemental Material for "The Neurodata Without Borders ecosystem for neurophysiological data science"

### Supplementary Material 1. Intracellular Electrophysiology Example using NWB and DANDI

The following shows a simple example, demonstrating the use of NWB for storage of intracellular electrophysiology data. The file used in this example is from DANDISET 20 (available at https://dandiarchive.org/dandiset/000020) from the Allen Institute for Brain Science as part of the multimodal characterization of cell types in the mouse visual cortex (see https://portal.brain-map.org/explore/classes/multimodal-characterization/multimodal-characterization-mouse-visual-cortex).

The following simple code example illustrates: 1) downloading of the file from DANDI, 2) reading the file with PyNWB, 3) visualization of the stimulus and response recording for a single sweep, and 4) visualization of the NWB file in NWB Widgets.

```
# import required libraries
from dandi.dandiapi import DandiAPIClient
from pynwb import NWBHDF5IO
from nwbwidgets import nwb2widget
from nwbwidgets.timeseries import show_indexed_timeseries_mpl
import numpy as np
from matplotlib import pyplot as plt

# Determine the s3path for the file on DANDI
dandiset_id = '000020'
filepath = 'sub-1001658946/sub-1001658946_ses-1003020741_icephys.nwb'
with DandiAPIClient() as client:
    asset = client.get_dandiset(dandiset_id, 'draft').get_asset_by_path(filepath)
    s3_path = asset.get_content_url(follow_redirects=1, strip_query=True)

# Open the file using the ros3 driver for streaming data access
nwb_s3io = NWBHDF5IO(s3_path, mode='r', load_namespaces=True, driver='ros3')
# Read the file from DANDI. Here we only read the structure and
# attributes of the file, but not the bulk data
nwbfile = nwb_s3f.read()

# Create a simple example visualization of the response and stimulus
# timeseries for a single sweep
# get the timeseries associated with a particular sweep number
sweep_number = 3
series = nwbfile.sweep_table.get_series(sweep_number)
# create a matplotlib figure for plotting
plt.rcParams['font.size'] = '16'
fig, (ax1, ax2) = plt.subplots(2, sharex=True, figsize=(12,8))
# plot the response and stimulus timeseries for the given sweep.
show_indexed_timeseries_mpl(series[0],
                            title=series[0].neurodata_type + " : " + series[0].name,
                            xlabel=None,
                            ax=ax1)
show_indexed_timeseries_mpl(series[1],
                            title=series[1].neurodata_type + " : " + series[1].name,
                            ax=ax2)
plt.show()
```

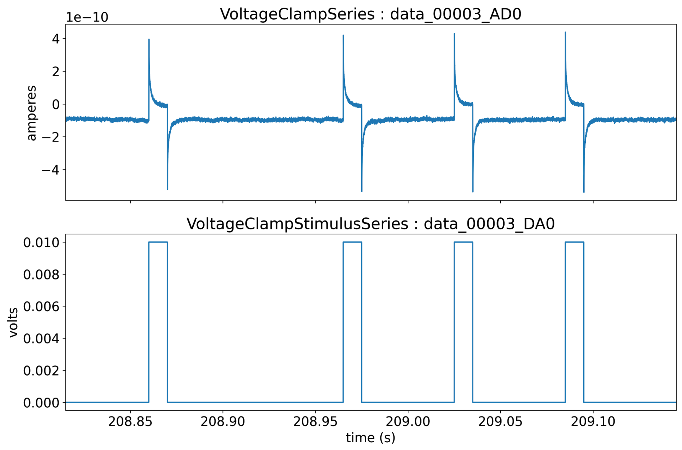

```
# Display the file with NWBWidgets
nwb2widget(nwbfile)
```

| | |
|---|---|
| session_description: | PLACEHOLDER |
| identifier: | b215c33e0839253a6c9ee5d6d0a6975cb12437110acc9a7c26f59374de8ee055 |
| session_start_time: | 2020-01-27 21:53:18.318000+00:00 |
| timestamps_reference_time: | 2020-01-27 21:53:18.318000+00:00 |
| session_id: | 1003020741 |
| institution: | Allen Institute for Brain Science |
| data_collection: | Free Memory: 14.6846 GB Please be patient while we export all existing acquired c… |
| source_script: | Release_2.0_20190602-977-g33b63e3b Date and time of last commit: 2020-05-29T… |
| source_script_file_name: | Vipr2-IRES2-Cre;Slc32a1-IRES2-FlpO;Ai65-508158.01.02.01 |

▾ **acquisition**

▸ **data_00000_AD0**

▸ **data_00001_AD0**

▸ **data_00002_AD0**

▾ **data_00003_AD0**

start (s) 208.82 duration (s): 0.3300

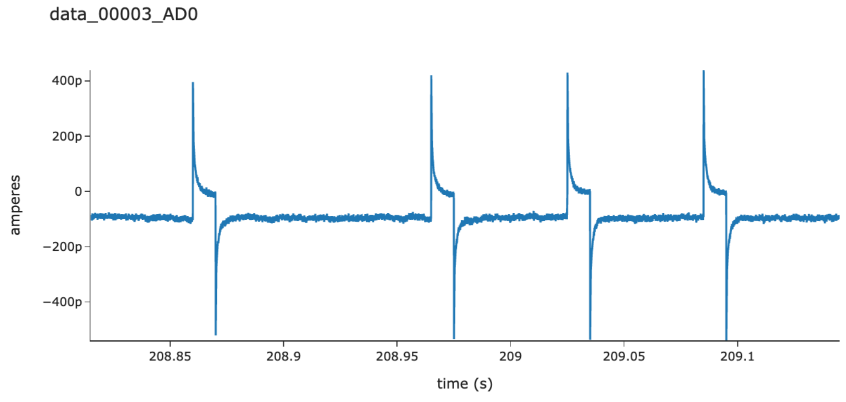

### Supplementary Material 2: Creating a New Extension

Using the ndx-simulation-output extension (see **Fig.4c**) as an example, we illustrate in the following the main steps for creating a new extension outlined in **Fig.4a2**.

#### 2.1 Set up new NDX using the NDX Template

The following code snippet shows the process for setting up the ndx-simulation-output extension using the ndx-template template. The template guides the developer through the setup via a simple question-and-answer process. In the code snippet, text shown in black is printed by cookiecutter, and text shown in blue are commands/responses entered by the developer.

```
>> cookiecutter gh:nwb-extensions/ndx-template
You've downloaded /Users/oruebel/.cookiecutters/ndx-template before. Is it okay to delete and
re-download it? [yes]: yes
namespace [ndx-my-namespace]: ndx-simulation-output
description [My NWB extension]: Data types for recording data from multiple compartments of
multiple neurons in a single TimeSeries.
author [My Name]: Ben Dichter
email []: ben.dichter@...
github_username [myname]: bendichter
copyright [2021, Ben Dichter]:
version [0.1.0]: 0.2.6
release [alpha]:
license [BSD 3-Clause]:
py_pkg_name [ndx_simulation_output]:
```

With these simple steps the template automatically sets up the full structure for our extension.

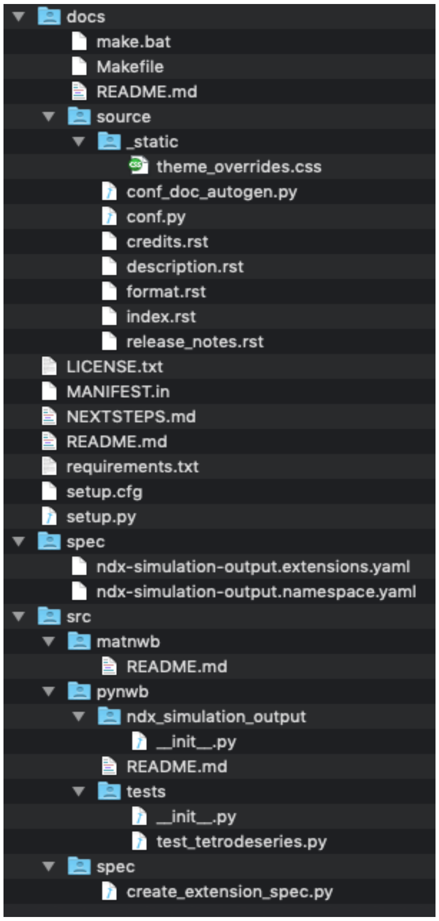

**Figure S2.1.** Files and folders generated by the cookiecutter ndx-template. The main folder contains the license and readme file for extension along with files required for installing the extension (e.g., *setup.py, setup.cfg, MANIFEST.in,* and *requirements.t*xt) as well a markdown file with instructions for next steps. The *docs/* folder contains the Sphinx documentation setup for the extension. Without any additional changes required, the developer can with this setup automatically generate documentation in HTML, PDF, ePub and many other formats directly from the extension schema using the HDMF-DocUtils. Generating the documentation is as simple as executing "*make html"* in the *docs/* folder. The *spec/* folder contains the schema files for the extensions. The schema files are generated by the script in */src/spec/create_extension_spec.py* (see **Sec. 2.2** next), and are typically not modified manually by the developers. The */src* folder then contains main source codes for the extension, including the: *spec/* folder with the code to generated the extension schema *matnwb/* folder with code for MatNWB *pynwb/* folder with code for PyNWB

### 2.2 Define the Extension Schema

The code example below shows the */src/spec/create_extension_spec.py* script to define and generate the schema for the *ndx-simulation-output* extension using the PyNWB data format specification API. Code shown in red has been auto-generated by the *ndx-template*. Code shown in blue has been defined by the developer to create the schema. Running this script then automatically generates the YAML schema files for the extension stored in the *spec/* folder.

```
# -*- coding: utf-8 -*-
import os.path
from pynwb.spec import NWBNamespaceBuilder, export_spec, NWBGroupSpec
def main():
    # these arguments were auto-generated from your cookiecutter inputs
    ns_builder = NWBNamespaceBuilder(doc='Data types for recording data from multiple compartments'
                                         'of multiple neurons in a single TimeSeries.',
                                     name='ndx-simulation-output',
                                     version='0.2.6',
                                     author='Ben Dichter',
                                     contact='')
    types_to_include = ['TimeSeries', 'VectorData', 'VectorIndex', 'DynamicTable', 'LabMetaData']
    for ndtype in types_to_include:
        ns_builder.include_type(ndtype, namespace='core')
    Compartments = NWBGroupSpec(default_name='compartments',
                                neurodata_type_def='Compartments',
                                neurodata_type_inc='DynamicTable',
                                doc='Table that holds information about '
                                    'what places are being recorded.')
    Compartments.add_dataset(name='number',
                             neurodata_type_inc='VectorData',
                             dtype='int',
                             doc='Cell compartment ids corresponding to a each column in the data.')
    Compartments.add_dataset(name='number_index',
                             neurodata_type_inc='VectorIndex',
                             doc='Index that maps cell to compartments.',
                             quantity='?')
    Compartments.add_dataset(name='position',
                             neurodata_type_inc='VectorData',
                             dtype='float',
                             quantity='?',
                             doc='Position of recording within a compartment. '
                                 '0 is close to soma, 1 is other end.')
    Compartments.add_dataset(name='position_index',
                             neurodata_type_inc='VectorIndex',
                             doc='Index for position.',
                             quantity='?')
    Compartments.add_dataset(name='label',
                             neurodata_type_inc='VectorData',
                             doc='Labels for compartments.',
                             dtype='text',
                             quantity='?')
    Compartments.add_dataset(name='label_index',
                             neurodata_type_inc='VectorIndex',
                             doc='indexes label',
                             quantity='?')
    CompartmentsSeries = NWBGroupSpec(neurodata_type_def='CompartmentSeries',
                                      neurodata_type_inc='TimeSeries',
                                      doc='Stores continuous data from cell compartments')
    CompartmentsSeries.add_link(name='compartments',
                                target_type='Compartments',
                                quantity='?',
                                doc='Metadata about compartments in this CompartmentSeries.')
    SimulationMetaData = NWBGroupSpec(name='simulation',
                                      neurodata_type_def='SimulationMetaData',
                                      neurodata_type_inc='LabMetaData',
                                      doc='Group that holds metadata for simulations.')
    SimulationMetaData.add_group(name='compartments',
                                 neurodata_type_inc='Compartments',
                                 doc='Table that holds information about '
                                     'what places are being recorded.')
    new_data_types = [Compartments, CompartmentsSeries, SimulationMetaData]
    # export the spec to yaml files in the spec folder
    output_dir = os.path.abspath(os.path.join(os.path.dirname(__file__), '..', '..', 'spec'))
    export_spec(ns_builder, new_data_types, output_dir)
if __name__ == "__main__":
    main()
```

#### 2.3 Create API Classes

The following code example is an abbreviated version of the *simulation_output.py* file located at *ndx-simulation-output/src/pynwb/ndx_simulation_output/* as part of the *ndx-simulation-output* extensions. For illustration purposes and to allow us to focus on the code relevant to the definition of the API classes, we here omit code details of the *find_compartments* function, which defines custom functionality.

The example shown here illustrates three main patterns for creating API classes for extensions. In part **A.** the extension uses the *get_class* method to dynamically generate an API Container class for the *SimulationMetaData* type directly from the schema. In part **B.,** the extension uses the same approach for *CompartmentSeries*, but then further customizes the class by adding the *find_compartments* to the class to provide additional user functionality. In part **C.** the extension then defines a custom API Container class for the *Compartments* type that extends the *DynamicTable* type.

```
# Import methods for registering and creating container class
from pynwb import register_class, docval, get_class
# Import the docval decorator used for documenting functions and type checking
from hdmf.utils import docval, call_docval_func
# Import the base Container classes we are extending
from hdmf.common.table import DynamicTable, ElementIdentifiers

# Define the name of the namespace of our extension needed to register Container classes
namespace = 'ndx-simulation-output'

# A. Auto-generate a Container class for the SimuluationMetaData type
SimulationMetaData = get_class('SimulationMetaData', namespace)

# B. Auto-generate a Container class for the CompartmentSeries type
CompartmentSeries = get_class('CompartmentSeries', namespace)
# B.1. Use monkey patching to add custom functionality to the auto-generated class
def find_compartments(self, cell, compartment_numbers=None, compartment_labels=None):
    [...]     # Details of the find_compartment omitted here for clarity.
CompartmentSeries.find_compartments = find_compartments

# C. Define a custom Container class for the Compartments table type
@register_class('Compartments', namespace) # Register the class with the TypeMap
class Compartments(DynamicTable):

    # Define the columns for the table. HDMF then automatically handles
    # setting up the columns for us as part of the class
    __columns__ = (
        {'name': 'number', 'index': True,
         'description': 'cell compartment ids corresponding to a each column in the data'},
        {'name': 'position', 'index': True,
         'description': 'the observation intervals for each unit'},
        {'name': 'label', 'description': 'the electrodes that each spike unit came from',
         'index': True, 'table': True}
    )

    # Document and define the allowable types for the parameters of the __init__ function
    @docval({'name': 'name', 'type': str,
             'doc': 'Name of this Compartments object','default': 'compartments'},
            {'name': 'id', 'type': ('array_data', ElementIdentifiers),
             'doc': 'the identifiers for the units stored in this interface', 'default': None},
            {'name': 'columns', 'type': (tuple, list),
             'doc': 'the columns in this table', 'default': None},
            {'name': 'colnames', 'type': 'array_data',
             'doc': 'the names of the columns in this table', 'default': None},
            {'name': 'description', 'type': str,
             'doc': 'a description of what is in this table',
             'default': 'Table that holds information about what places are being recorded.'},
            )
    def __init__(self, **kwargs):
        call_docval_func(super(Compartments, self).__init__, kwargs)
```

#### 2.4 Documenting the Extension

The *ndx-template* automatically generates as part of the *docs/* folder the full setup for automatically generating Sphinx-based documentation for the extension from the schema using the hdmf-docutils library. To generate the documentation we simply need to run the command *"make html"* in the *docs/* folder. Using the same approach we can generate documentation in a large range of common formats, e.g., HTML, PDF, man, or ePub. The ndx-template also generates standard *credits.rst*, *format.rst*, *release_notes.rst*, *description.rst*, and *index.rst* source files to make it easy for developers to customize the documentation and include additional details about the extension.

### Supplementary Material 3. Process for Creating, Publishing, and Updating Neurodata Extensions (NDX)

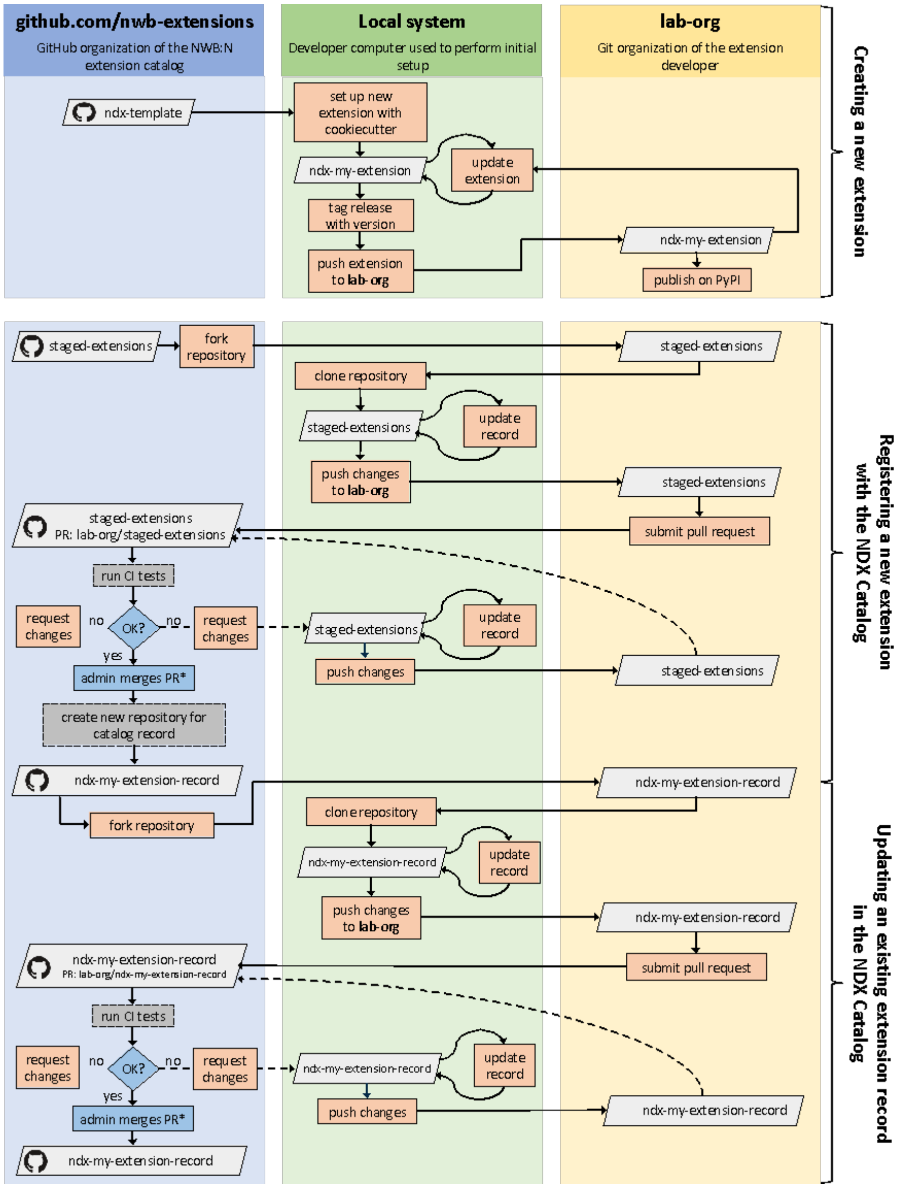

**Figure S3.1** Illustration of the process for creating, publishing, and updating extensions via the Neurodata Extension Catalog (NDX Catalog), and (3) updating an extension/record. Boxes shown in gray indicate Git repositories; boxes in orange describe user actions; and boxes in blue indicate actions by administrators of the NDX catalog.

**Figure S3.1** shows an overview of the process for (1) creating a new extension, (2) creating a record to publish an extension via the Neurodata Extension Catalog (NDX Catalog), and (3) updating an extension/record. The figure also illustrates the automated CI processes that are managed in the NDX Catalog. The catalog process is modeled after the conda-forge model, which enables automation of many catalog processes using free, public services and avoiding the need for NWB to host its own services.

In the NDX Catalog, extensions are shared via the dedicated *nwb-extensions* GitHub organization for the NDX Catalog (blue column). The NDX Catalog provides the *ndx-template* cookiecutter extension template repository as well as the *staged-extensions* repository for submitting extensions to the NDX Catalog. Each extension record is then managed in a corresponding ndx record Git repository as part the *nwb-extensions* GitHub organization. The public NDX record repositories contains a *README.md* file describing the extension along with a *ndx-meta.yaml* metadata record for extensions, with basic information required for installing and locating the extension (see **Fig. S3.2**).

Creation and changes to the extension on record are usually performed by a developer on their local system, e.g., laptop computer (green column). The developer then submits the changes to the extension or record repository via pull requests.

In this process, the Git repository with the sources of the extension remains in the lab organization of the submitter (yellow column). Here the only requirements are that: 1) the extension is stored in a Git repository and 2) the repository must be publicly accessible via the organization of the submitter, such that the repository can be cloned directly from the source location indicated in the NDX metadata record (**Fig. S3.2**). This strategy allows for labs, universities, and independent groups to maintain ownership of the source code for their extensions in their own public Git space (e.g, on GitHub, Bitbucket, or GitLab) while creating an open, standardized record of all public extensions in a central location as part of the NDX Catalog. The ability for submitters to retain ownership of their extensions in their own organization is important to facilitate development as well as to retain a clear chain of responsibility and ownership. This is particularly important when the developers of the extension are funded by their own grants and/or are applying for funding.

```yaml
name: ndx-simulation-output
version: 0.2.6
src: https://github.com/bendichter/ndx-simulation-output
pip: https://pypi.org/project/ndx-simulation-output
license: BSD
maintainers:
  - bendichter
```

**Figure S3.2.** Example *ndx-meta.yaml* metadata record for the *ndx-simulation-output* extension.

### Supplementary Material 4: Overview of Select Data Analysis, Visualization, and Management Tools That Support NWB

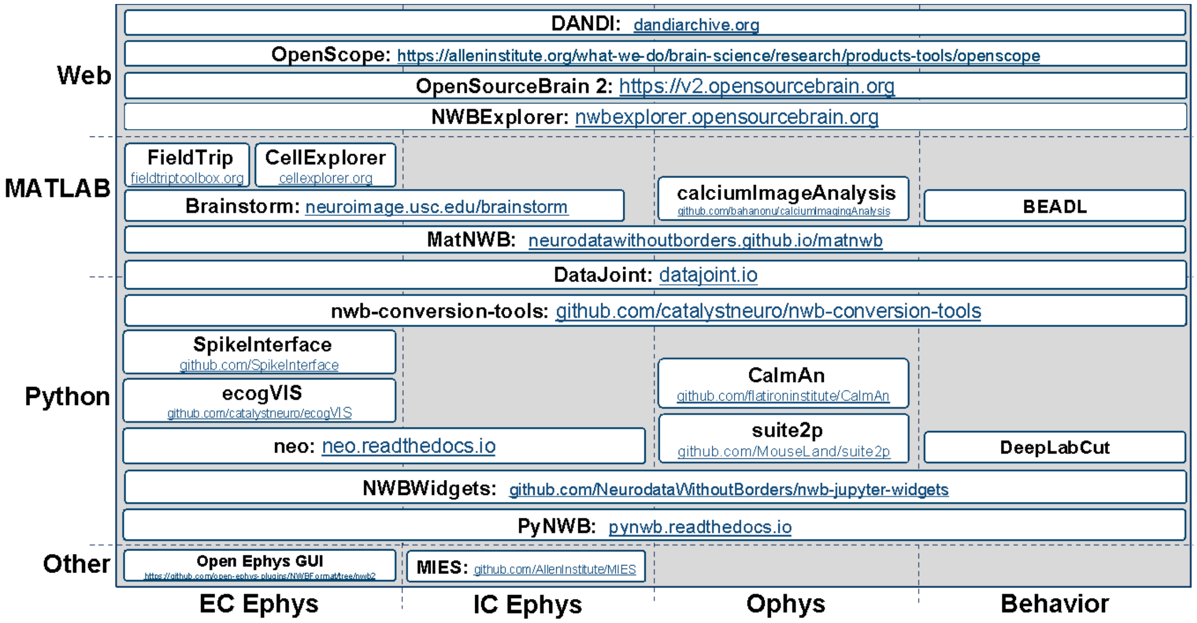

**Figure S.4.** Visualization showing select data analysis, visualization, and management tools that support NWB organized by their main application (x-axis) and programming environment (y-axis).

### Supplementary Material 5. Assessment of FAIRness of NWB + DANDI

Table 5.1 – 5.4 below assess different solutions for sharing neurophysiology data with regard to their compliance with FAIR data principles, with cells shown in: **i)** gray indicate non-compliance, **ii)** green indicate compliance, and **iii)** yellow indicate partial compliance either due to incomplete implementation or optional support, leaving achieving compliance ultimately to the end user. The assessment for NIX is based on the INCF review for SPP endorsement[1]. The *"Custom"* row in the tables refers to lab-specific binary formats.

In practice, the various approaches target different principle uses, and as such this is not an assessment of the quality of a product per-se, but rather its out-of-the-box compliance with FAIR principles in the context of neurophysiology. For example, self-describing data formats (like HDF5 or Zarr) seek to address challenges in high-performance data management and storage independent of a particular application, and as such lack specifics about (meta)data related to neurophysiology. However, while self-describing formats (like HDF5) are by themselves not sufficient to achieve FAIR compliance, they still form a critical building block in an overall strategy for FAIR data as evidenced by the fact that NIX, NWB, and many other application standards across science domains build on HDF5. Similarly, NIX provides a generic data model to enable storage of *"fully annotated scientific datasets, i.e. the data together with its metadata within the same container*" with the goal to enable "*standardization by providing a common/generic data structure for a multitude of data types."*[2] As such, NIX provides important functionality towards building a FAIR data strategy, but the NIX data model by itself lacks specificity with regard to neurophysiology, leaving it up to the user to define appropriate schema to facilitate FAIR compliance. Broadly speaking, with increasing specificity of data standards——i.e., as we move from general-purpose, self-describing formats (Zarr, HDF5) to generic data standards (NIX) to application-specific standards (NWB)— compliance with FAIR principles and rigidness of the data specification increases.

[1] M. Martone. R. Gerkin, R. Moucek, S. Das, W. Goscinski, J. Hellgren-Kotaleski, D. Kennedy, T.Leergaard, J. Boline, M. Abrams, "SBP Review: NIXV1.0, " May 13, 2020, DOI: https://doi.org/10.7490/f1000research.1117858.1

[2] Text in italic quoted from http://g-node.github.io/nix/

| | Findable | | | |
|---|---|---|---|---|
| | F1. (Meta)data are assigned a globally unique and persistent identifier | F2. Data are described with rich metadata (defined by R1 below) | F3. Metadata clearly and explicitly include the identifier of the data they describe | F4. (Meta)data are registered or indexed in a searchable resource |
| Custom | No | No | No | • N/A. This is a key function of data archives and management systems |
| Zarr | No | • Self-describing, structural metadata (e.g., data type, array shape etc.) only<br>• Scientific (meta)data is fully user defined | | |
| HDF5 | No | | | |
| NIX | • UUIDs are assigned to all objects | • Self-describing, structural metadata (uses HDF5)<br>• Generic data model (i.e., scientific (meta)data is user-defined) | | |
| NWB 1.0 | No | • Yes, but the schema language was not formally defined | • Similar to NWB 2.x but the much more flexible schema (including inclusion of arbitrary data) often lead to non-compliance | |
| NWB 2.x | • UUIDs are assigned to all objects<br>• External file identifier can be stored in the *identifier* field | • Rich schema for neurophysiology (meta)data<br>• Self-describing, structural metadata (uses HDF5) constrained by the standard schema | • Metadata is either directly associated with or explicitly linked to by the corresponding objects | |
| DANDI | • All dandisets and assets carry unique and persistent identifiers | • Uses NWB and other modern data standards<br>• Provides its own Dandiset schema for metadata about whole data collections | • Yes, persistent identifiers used by the archive are included with the metadata | • DANDI is a public archive that features rich search features over publicly shared data |

**Table 5.1** Compliance of NWB+DANDI with FAIR principles: Findability

<table>
<tr><th></th><th colspan="4">Accessible</th></tr>
<tr><th></th><th>A1. (Meta)data are retrievable by their identifier using a standardised communications protocol</th><th>A1.1 The protocol is open, free, and universally implementable</th><th>A1.2 The protocol allows for an authentication and authorisation procedure, where necessary</th><th>A2. Metadata are accessible, even when the data are no longer available</th></tr>
<tr><td>Custom</td><td>No</td><td>No</td><td rowspan="6">• N/A. This is a key function of data archives and management systems<br>• Encryption of files is possible via external tools<br>• HDF5/Zarr could support encryption of data elements via I/O filters</td><td rowspan="6">• N/A. This is a key function of data archives and management systems</td></tr>
<tr><td>Zarr</td><td rowspan="2">• Non-persistent file/object paths only</td><td>• Yes, but python-only API<br>• Long-term support is not clear</td></tr>
<tr><td>HDF5</td><td>• Portable format with broad support across programming languages and compute systems<br>• Intended for long-term support</td></tr>
<tr><td>NIX</td><td>• Yes</td><td>• Uses HDF5<br>• NIX API for C++. Matlab, Python and Java<br>• Open source</td></tr>
<tr><td>NWB 1.0</td><td>• Non-persistent file/object paths only (same as HDF5)</td><td>• Yes, but schema language was not formally defined and available APIs were limited</td></tr>
<tr><td>NWB 2.x</td><td>• Yes. Objects retrievable based on UUID and path.</td><td>• Uses HDF5<br>• NWB API in Python and Matlab<br>• Open source</td></tr>
<tr><td>DANDI</td><td>• Uses NWB<br>• Metadata is exported as JSON/JSON-LD alongside with data</td><td>• Uses standard protocols (e.g., REST API)</td><td>• Supports user authentication and authorized access to all Dandisets,</td><td>• Searchable on the the archive and exposed as LinkedData</td></tr>
</table>

| | | | |
|---|---|---|---|
| | • REST API, Python, CLI, DataLad, ROS3 HDF5 | • Supports integration with external services | assets and other DANDI resources |

**Table 5.2** Compliance of NWB+DANDI with FAIR principles: Accessibility

| | **Interoperable** | | |
|---|---|---|---|
| | I1. (Meta)data uses a formal, accessible, shared, and broadly applicable language for knowledge representation. | I2. (Meta)data use vocabularies that follow FAIR principles | I3. (Meta)data include qualified references to other (meta)data |
| Custom | No | No | No |
| Zarr | No | No | No |
| HDF5 | No | No | No |
| NIX | • Uses odML<br>• Uses HDF5 | • User defined | • User defined |
| NWB 1.0 | • Uses custom schema definition in Python | • Data follows the NWB 1.0 schema | • Partially. NWB 2.x significantly enhanced support for linking of metadata with data. |
| NWB 2.x | • Schema defined in JSON/YAML using json-schema<br>• NWB and extension schema are available with NWB files and online<br>• Uses HDF5 | • Data follows the NWB schema<br>• NWB supports use of ontologies via linking to external resources[3] | • The NWB schema explicitly models links between (meta)data<br>• NWB supports linking to external resources[3] |
| DANDI | • Uses NWB, JSON + json-schema, JSON-LD | • Uses NWB and other FAIR ontologies | • schema.org, spdx.org (licenses), PROV |

**Table 5.3** Compliance of NWB+DANDI with FAIR principles: Interoperability

[3] Support for external resources has been released in HDMF >2.3 and is currently undergoing community review for integration with the NWB core data standard.

| | Reusable | | | |
|---|---|---|---|---|
| | R1. (Meta)data are richly described with a plurality of accurate and relevant attributes | R1.1. (Meta)data are released with a clear and accessible data usage license | R1.2. (Meta)data are associated with detailed provenance | R1.3. (Meta)data meet domain-relevant community standards |
| Custom | No | • N/A. Usage licences are typically managed by data archives | No | No |
| Zarr | No | | No | No |
| HDF5 | No | | No | No |
| NIX | • User defined | | No | • User defined |
| NWB 1.0 | • Yes | | • Yes. NWB 2.x further refined this significantly | • Yes |
| NWB 2.x | • Yes | | • Includes detailed metadata about publications, experimenters, devices, subjects etc.<br>• Derived data (e.g., ROIs) link to the source data | • Yes, NWB provides detailed, neurophysiology-specific data schema |
| DANDI | • Uses NWB and defined dandiset schema | • All data in DANDI is published with a clear data usage licence | • Dandisets support detailed metadata about the data generation<br>• Dandisets are versioned | • Uses NWB |

**Table 5.4** Compliance of NWB+DANDI with FAIR principles: Reusability

### Supplementary Material 6. Diverse Community of Data Producers Adopting NWB.

The table shown below provides an overview of select labs that are using NWB. The last column of the table lists relevant DANDI datasets that have been published via the DANDI data archive using NWB. Each DANDI dataset typically consists of a large collection of NWB files related to a particular publication or experiment, with each NWB file representing the data from a particular recording session. All DANDI datasets can be found online at https://dandiarchive.org/dandiset/{6-digit-zero-padded-id}, e.g., https://dandiarchive.org/dandiset/000007. In the "Modality" column of the table we use the following abbreviations:

- ecephys: extracellular electrophysiology
- icephys: intracellular electrophysiology
- ophys: optical physiology
- ECoG: Electrocorticography
- fNIRS: Functional near-infrared spectroscopy

| Name, Affiliation | Species | Modality | DANDI datasets |
|---|---|---|---|
| AE Studio | human | fNIRS | 122 |
| Allen Institute | mouse, human | ecephys, icephys, ophys | 12, 20, 21, 22, 23, 24, 30, 36, 37, 39, 42, 43, 48, 49, 50, 66, 107, 109, 142, 209 |
| R. Axel, Columbia | fly | ophys | |
| Blue Brain Project | mouse | icephys | 25 |
| J. Berke, UCSF | rat | ecephys | |
| K. Bouchard, LBNL/UC Berkeley | rat, simulation | ecephys, uECoG | |
| C. Brody, Princeton | rat, mouse | ecephys | |
| B. Brunton, U Washington | human | ECoG | 55 |
| E. Buffalo, U Washington | monkey | ecephys | |
| T. Buschman, Princeton | monkey | ecephys | |
| G. Buzsaki, NYU | rat, mouse | ecephys | 3, 41, 44, 56, 59, 61, 67, 166, 213, 218 |
| M. Capogna, Aarhus | mouse | ecephys | |
| M. Carandini, UCL | mouse | ecephys | 17 |
| E. Chang, UCSF | human | ECoG | 19 |

| | | | |
|---|---|---|---|
| A. Churchland, CSHL | mouse | ecephys, ophys | 16 |
| R. Cossart, Inserm | mouse | ophys | 219 |
| D. Feldman, UC Berkeley | mouse | ecephys | |
| A. Fleischmann, Brown | mouse | ophys | 167 |
| L. Frank, UCSF | mouse | ecephys | 65, 115, 165 |
| L. Giocomo, Stanford | mouse | ecephys, ophys | 53, 54 |
| A. Groh, Heidelberg | mouse | ecephys, ophys | |
| K. Harris, UCL | mouse | ecephys | 17 |
| M. Hennig, Edinburgh | mouse | ecephys | 28, 34 |
| S. Husainni, Columbia | mouse | ecephys | |
| International Brain Lab | mouse | ecephys | 45, 149 |
| M. Jazayeri, MIT | monkey | ecephys | 130 |
| D. Jaeger, Emory | mouse | ophys, ecephys, icephys | |
| S. Kastner, Princeton | monkey | ecephys | |
| N. Li, Baylor | mouse | ecephys | 7 |
| A. Losonczy, Columbia | mouse | ophys | |
| G. Maimon, Rockefeller | fly | behavior | 212 |
| J. Martinez, Western | mouse, monkey, human | icephys | |
| R. McGreal, UCSB | mouse | ophys | 206 |
| L. Miller, Northwestern | monkey | ecephys | 127 |
| T. Movshon, NYU | monkey | ecephys | |
| D. O'Connor, Johns Hopkins | mouse | ecephys, ophys | |
| J. Parvizi, Stanford | human | ECoG | |
| U. Rutishauser, Cedars-Sinai | human | ecephys | 4, 207 |
| B. Sabatini, Harvard | mouse | icephys | |
| S. Schultz, Imperial | mouse | ecephys, ophys | |
| K. Shenoy, Stanford | monkey | ecephys | 70, 121 |
| M. Smear, U Oregon | mouse | behavior | 217 |

| | | | |
|---|---|---|---|
| S. Smith, UCSB | mouse | ophys | 206 |
| I. Soltesz, Stanford | mouse, simulation | ophys, icephys | |
| N. Steinmetz, U Washington | mouse | ecephys | 17 |
| K. Svoboda, Janelia | mouse | ecephys, ophys | 5, 6, 9, 10, 11, 13, 15, 60, 168 |
| N. Tandon, UT Houston | human | ECoG | |
| D. Tank, Princeton | mouse | ecephys | |
| H. Tao, USC | mouse | icephys | 117 |
| A. Tolias, Baylor | mouse | icephys | 8, 35 |
| S. Tripathy, UofToronto/CAMH | human, mouse | icephys | |
| T. Valiante, Toronto | human | icephys | |

### Supplementary Material 7: Software Release Process and History

Software releases and processes are one indicator for the maturity of software products. As such, looking at the release history of NWB also provides some insight into how NWB has evolved over the course of the project from a first prototype of NWB 2 to a production data standard and software ecosystem.

The initial development of NWB 2 occurred during Nov. 2016 – Nov.2017. This phase did not include formal releases as the focus was on agile and rapid development of functionality with the goal to establish design principles and create a usable, fully functional prototype. Changes to the standard and software were evaluated in this phase by early adopters and reviewed by the community as part of NWB community hackathons.

The first beta release of the NWB 2 schema, PyNWB 0.2.0, and MatNWB 0.1.0b then occurred in November 2017 in conjunction with SfN. This marked the start of the beta testing and development phase of NWB, which occurred between Nov.2017 – Jan.2019. During the beta phase NWB adopted a more formal release process of versioned releases via pip, conda, and GitHub. These releases were targeted at early adopters and beta testers while development was still largely agile based on GitHub source releases. During this phase the focus was on evaluation, refinement, and productization.

In Jan.2019 we then released the first official version of NWB 2, including nwb-schema 2.0, PyNWB 1.0, and matnwb 0.2.0. This marked the beginning of the adoption and integration phase for the NWB 2 project. During Jan.2019 – Apr. 2021, a key focus has been on the one hand to continue to advance and refine NWB to meet the needs of adopters as well as to work with neuroscience labs and tool developers to support NWB. With the shift in focus from development to adoption then also came further refinement of the release processes and adoption of stricter software versioning guidelines based on semantic versioning to facilitate integration of NWB software with other software tools and adoption in lap data pipelines. While software releases in this phase were still often determined on a per-need-basis, the goal was to keep the APIs and standard as stable as possible.

One strategy to achieve this goal then was to separate the core data modeling capabilities from PyNWB into the separate HDMF library. Publishing HDMF as its own software product has been essential both to facilitate reuse of HDMF capabilities for other applications as well as to ensure stability of the PyNWB user API. As shown in **Fig. S.7**, PyNWB has undergone only 1 major release and 5 minor releases since the first release of PyNWB 1.0.0. At the same time, HDMF underwent a much larger number of releases. This illustrates the effectiveness of the approach of separating core infrastructure from user-APIs, as it allowed us to continue to advance core NWB technologies while limiting impact on end users. Similarly, extracting general schema (e.g., for dynamic data tables) into the separate hdmf-common-schema allowed to further make these common building blocks broadly accessible to science applications and to continue to develop them as part of the HDMF core software infrastructure.

MatNWB, through its strategy to auto-generate API classes directly from the NWB schema, is tied directly in a particular release to the most recent version of the NWB schema that the particular release supports. In April 2020, MatNWB, therefore, adopted a new, extended semantic versioning scheme for its software release consisting of 4 digits, with the first 3 digits indicating the major, minor, and patch release of the NWB schema and the last digit indicating the software patch release of MatNWB. As such, version 2.2.5.1 of MatNWB supports NWB schema 2.2.5 as the most recent version of the NWB format and includes 1 software patch release of MatNWB.

MatNWB then also added all tagged versions of the NWB schema with each release to avoid the need for git checkouts during the use of the software and facilitate interaction with NWB files with varying schema versions.

With the start of the NIH U24 project in April 2021, NWB then entered its next main phase with a focus on more widespread adoption to advance standardization of neurophysiology data through dissemination and integrating of NWB. With this transition, then also comes the need for further refinement of software release processes. This also means that adoption and integration projects increasingly no longer involve the NWB team directly, but are being led independently by other project teams. To facilitate planning and interaction, this required further refinement of release processes to adopt more rigid release plans and schedules and to facilitate contribution of other projects to NWB with predictable release timelines.

In addition to the software, key components are also releases of the NWB schema. Here, a main goal has been stability to ensure that files remain accessible. This is also reflected in the release history of the NWB schema, which has undergone only four minor releases and no major releases since the first full release of the schema. These releases largely focused on addition and refinement of data schema, while the APIs support reading of data of all NWB 2.x file versions.

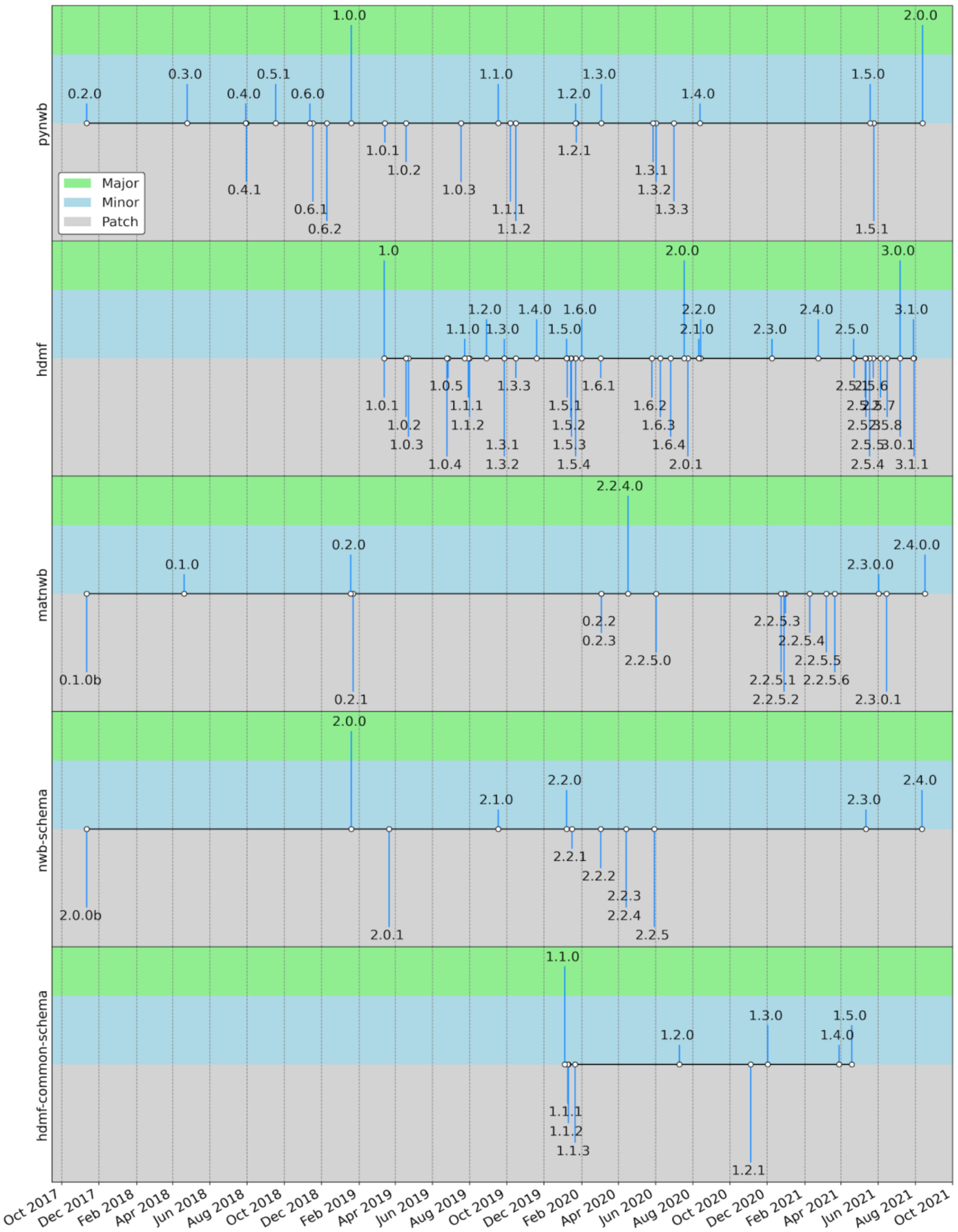

**Figure S.7.** Overview of the release history of the PyNWB, HDMF, and MatNWB APIs and the NWB and hdmf-common data standard schema.

### Supplementary Material 8. NWB Online and Social Media Resources

NWB online and social media resources provide additional resources for users and the broader community to engage with and learn about NWB. The nwb.org website[69] serves as the central entry-point for users to NWB and provides high-level information about NWB and links to all relevant online resources and tools discussed in the Methods. Additional online resources include **[Slack]**[70]**, [Twitter]**[71]**, [YouTube]**[72], and the **[NWB Mailing List]**[73].

### Supplementary Material 9. NWB Training Resources

NWB provides a broad range of training resources for users and developers. Users who want to learn more about how to use NWB can view the NWB online video training course as part of the **[INCF Training Space]**[74]. Detailed code tutorials are further available as part of the **[PyNWB Documentation]**[58] and **[MatNWB Documentation]**[60].

For Neurodata Extensions (NDX), detailed documentation of versioning guidelines, sharing guidelines and strategies, and the proposal review process are available online as part of the NDX Catalog[66]. Step-by-step instructions for creating new NDX are provided as part of the **[NWB Extensions Template]**[67].

Additional resources for developers and data managers include the API documentation for **[HDMF]**[51]**, [PyNWB]**[58], and **[MatNWB]**[60] and documentation of the format schema as part of the **[NWB Schema]**[63], **[HDMF Common Schema]**[55]**, [NWB Storage]**[65]**,** and **[Specification Language]**[64].
